# The ENCODE Uniform Analysis Pipelines

**DOI:** 10.1101/2023.04.04.535623

**Authors:** Benjamin C. Hitz, Jin-Wook Lee, Otto Jolanki, Meenakshi S. Kagda, Keenan Graham, Paul Sud, Idan Gabdank, J. Seth Strattan, Cricket A. Sloan, Timothy Dreszer, Laurence D. Rowe, Nikhil R. Podduturi, Venkat S. Malladi, Esther T. Chan, Jean M. Davidson, Marcus Ho, Stuart Miyasato, Matt Simison, Forrest Tanaka, Yunhai Luo, Ian Whaling, Eurie L. Hong, Brian T. Lee, Richard Sandstrom, Eric Rynes, Jemma Nelson, Andrew Nishida, Alyssa Ingersoll, Michael Buckley, Mark Frerker, Daniel S Kim, Nathan Boley, Diane Trout, Alex Dobin, Sorena Rahmanian, Dana Wyman, Gabriela Balderrama-Gutierrez, Fairlie Reese, Neva C. Durand, Olga Dudchenko, David Weisz, Suhas S. P. Rao, Alyssa Blackburn, Dimos Gkountaroulis, Mahdi Sadr, Moshe Olshansky, Yossi Eliaz, Dat Nguyen, Ivan Bochkov, Muhammad Saad Shamim, Ragini Mahajan, Erez Aiden, Tom Gingeras, Simon Heath, Martin Hirst, W. James Kent, Anshul Kundaje, Ali Mortazavi, Barbara Wold, J. Michael Cherry

## Abstract

The Encyclopedia of DNA elements (ENCODE) project is a collaborative effort to create a comprehensive catalog of functional elements in the human genome. The current database comprises more than 19000 functional genomics experiments across more than 1000 cell lines and tissues using a wide array of experimental techniques to study the chromatin structure, regulatory and transcriptional landscape of the *Homo sapiens* and *Mus musculus* genomes. All experimental data, metadata, and associated computational analyses created by the ENCODE consortium are submitted to the Data Coordination Center (DCC) for validation, tracking, storage, and distribution to community resources and the scientific community. The ENCODE project has engineered and distributed uniform processing pipelines in order to promote data provenance and reproducibility as well as allow interoperability between genomic resources and other consortia. All data files, reference genome versions, software versions, and parameters used by the pipelines are captured and available *via* the ENCODE Portal. The pipeline code, developed using Docker and Workflow Description Language (WDL; https://openwdl.org/) is publicly available in GitHub, with images available on Dockerhub (https://hub.docker.com), enabling access to a diverse range of biomedical researchers. ENCODE pipelines maintained and used by the DCC can be installed to run on personal computers, local HPC clusters, or in cloud computing environments *via* Cromwell. Access to the pipelines and data *via* the cloud allows small labs the ability to use the data or software without access to institutional compute clusters. Standardization of the computational methodologies for analysis and quality control leads to comparable results from different ENCODE collections - a prerequisite for successful integrative analyses.

**Database URL:** https://www.encodeproject.org/

## Introduction

The Encyclopedia of DNA Elements (ENCODE) project^1^ (https://www.encodeproject.org/; Kagda et al, in preparation) is an international consortium with a goal of annotating regions of the human and mouse genomes. ENCODE aims to identify functional elements by investigating DNA and RNA binding proteins, chromatin structure, transcriptional activity and DNA methylation states for different biological samples. During the third and fourth phases of ENCODE (2012-2022) the diversity and volume of data increased as new genomic assays were added to the project. The diversity of biological samples used in these investigations has been expanded, including data from additional species (*D. melanogaster* and *C. elegans via* our sister projects modENCODE^2^; http://www.modencode.org) and experimental data are validated and analyzed using new methods. During the first 6 years of the pilot and initial scale-up phase, the project surveyed the landscape of the *H. sapiens* and *M. musculus* genomes using over 20 high-throughput genomic assays in more than 350 different cell and tissue types, resulting in over 3000 datasets. In addition to ENCODE funded projects, the DCC also has incorporated over 2000 datasets from the Roadmap for Epigenomics Consortium^3^ (REMC), The Genomics of Gene Regulation project (GGR; https://www.genome.gov/Funded-Programs-Projects/Genomics-of-Gene-Regulation), and the genomics community. The Data Coordination Center (DCC) is entrusted with validating, tracking, storing, visualizing, and distributing these data files and their metadata to the scientific community.

Uniform pipelines, the series of software algorithms that process raw sequencing data and generate interpretable data files, are important for scientific reproducibility. Publicly available pipelines allow researchers conducting similar experiments to share pipelines directly, making the results uniform and comparable. Multiple analysis pipelines exist for many assays and often differ in the software used for each component, the parameters defined for these components, or the statistical analysis used to determine significance of the results. The results from different pipelines for a given assay cannot always be appropriately compared. Thus, it is imperative for integrative analysis that results have the same basic assumptions, such as what defines a binding site, what reference genome is used, annotation standards for RNAs, cutoff used to define significance, etc. Historically, it has required significant technical expertise to set up, maintain, and run a single genomics analysis pipeline on local hardware. The ENCODE corpus contains over 80,000 fastq files across over 17,000 functional genomics experiments, with the majority being ChIP-Seq^4^, RNA-Seq^5^, or DNase-Seq^6^. The ChIP-seq pipeline works on both traditional transcription factors with narrow peak sizes, and histone mark ChIP experiments with broader peaks. The pipeline has been further modified for the multiplexed MINT-Chip^7^ assays. These pipelines were originally described^8^ but have been continuously modified as the ENCODE project has progressed and more data has been analyzed. We have implemented five RNA-Seq pipelines: One for typical transcripts (size selected at >200bp), one for shorter transcripts (size selected at <200bp), RAMPAGE^9^ and CAGE^10^, one for long-read RNA-seq, and one for micro-RNA-seq. We have also implemented pipelines for DNase-seq, ATAC-seq, Hi-C, and Whole-Genome Bisulfite Sequencing (WBGS). Help, descriptions, and ENCODE data standards can be found on the ENCODE Portal: https://www.encodeproject.org/pages/pipelines.

A bioinformatics analysis pipeline can be described as a series of computational steps, with defined (typically file-based) inputs and outputs, along with a set of parameters. The outputs of earlier steps in the pipeline are the inputs for later steps. Each “step”, which may be composed of one or more pieces of software, can be containerized in a system such as Docker (https://www.docker.com) to allow rapid and flexible provisioning of virtual computer systems to run the calculation specified. A typical genomics experiment has two or three major steps and may have other additional steps (although when replicate concordance calculations are involved, the process can get significantly more complicated). In a typical genomics pipeline, raw sequence data in the form of fastq files is mapped to the specific reference genome to produce one or more alignment files in BAM format^11^. The BAM files are then processed into one or more signal (typically bigWig^12^) and interval or “peak” files (bed and bigBed^12^). RNA-seq analysis typically has a transcript quantification step instead of peak calling, and produces a tab-delimited (tsv) file representing the expression for each gene or transcript. In addition to these “core” steps, the pipeline may require additional steps such as filtering, quality control metric calculations, and file format conversions.

These steps are defined and linked together using the Workflow Description Language^13^ (WDL), a domain-specific language developed at the Broad Institute. The WDL file defines each step, registers the input and output files and parameters, and provisions the resources as needed. With the onset of the fourth and final phase of the ENCODE project, we aspired to provide pipelines that could be run on a wide variety of platforms, either in the cloud or on local HPC systems. To this end, we adopted the Cromwell^14^ framework to manage execution of the pipeline code, input and output files across a variety of platforms including Google Cloud, Amazon Web Services, and local compute clusters using both Docker and Singularity (Fig 1).

**Figure 1.**
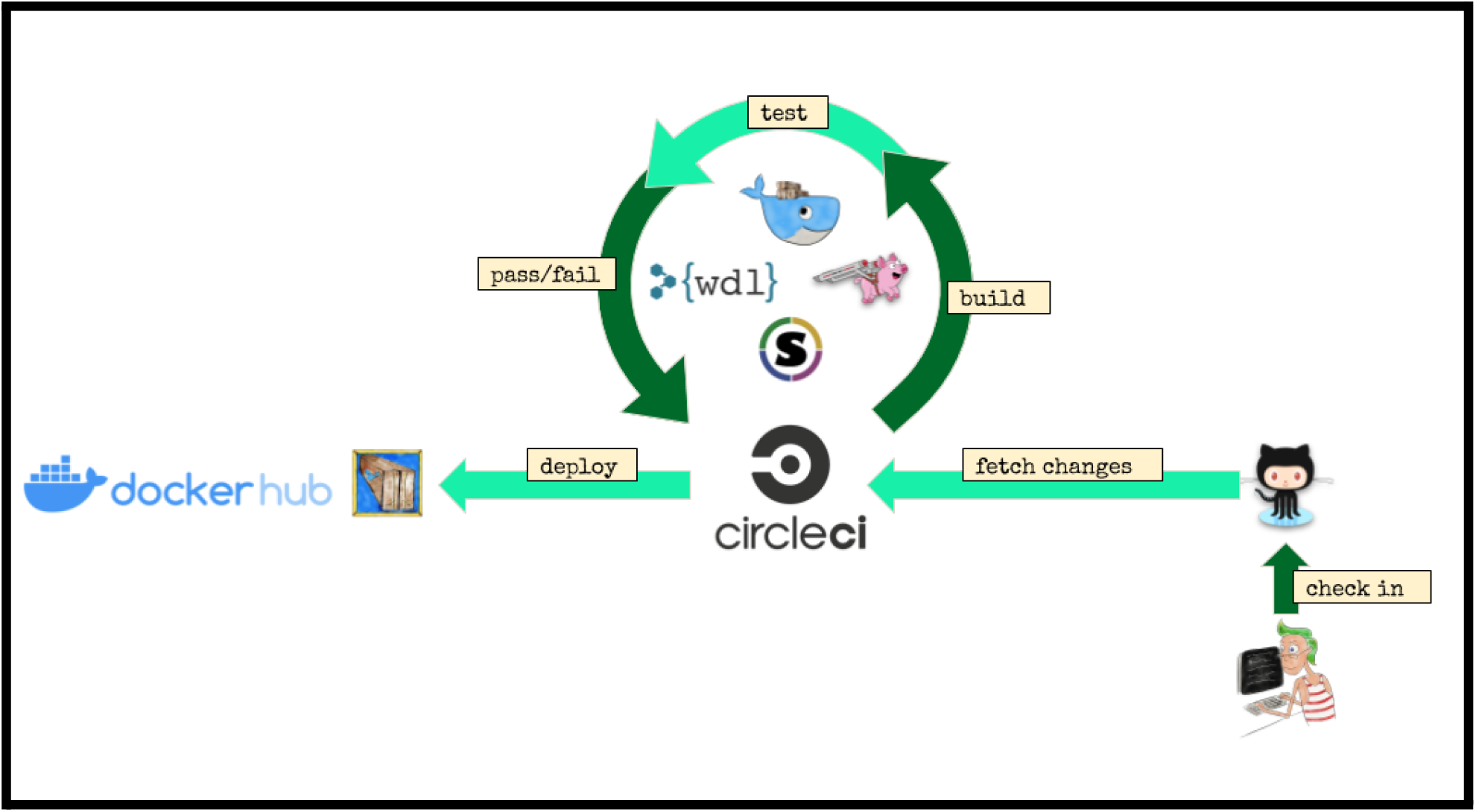
Pipeline infrastructure and continuous integration.

The code for all of the ENCODE pipelines use a common template, so the knowledge and understanding of the framework around one ENCODE pipeline is applicable to all the others. We have implemented unit testing, step-wise and end-to-end testing using circle-ci (https://www.circleci.com) for continuous integration, testing, and automatic docker builds. All code is available on GitHub and supported *via* GitHub issues. An example “demo” WDL pipeline is shown in Figure 2A.

**Figure 2.**
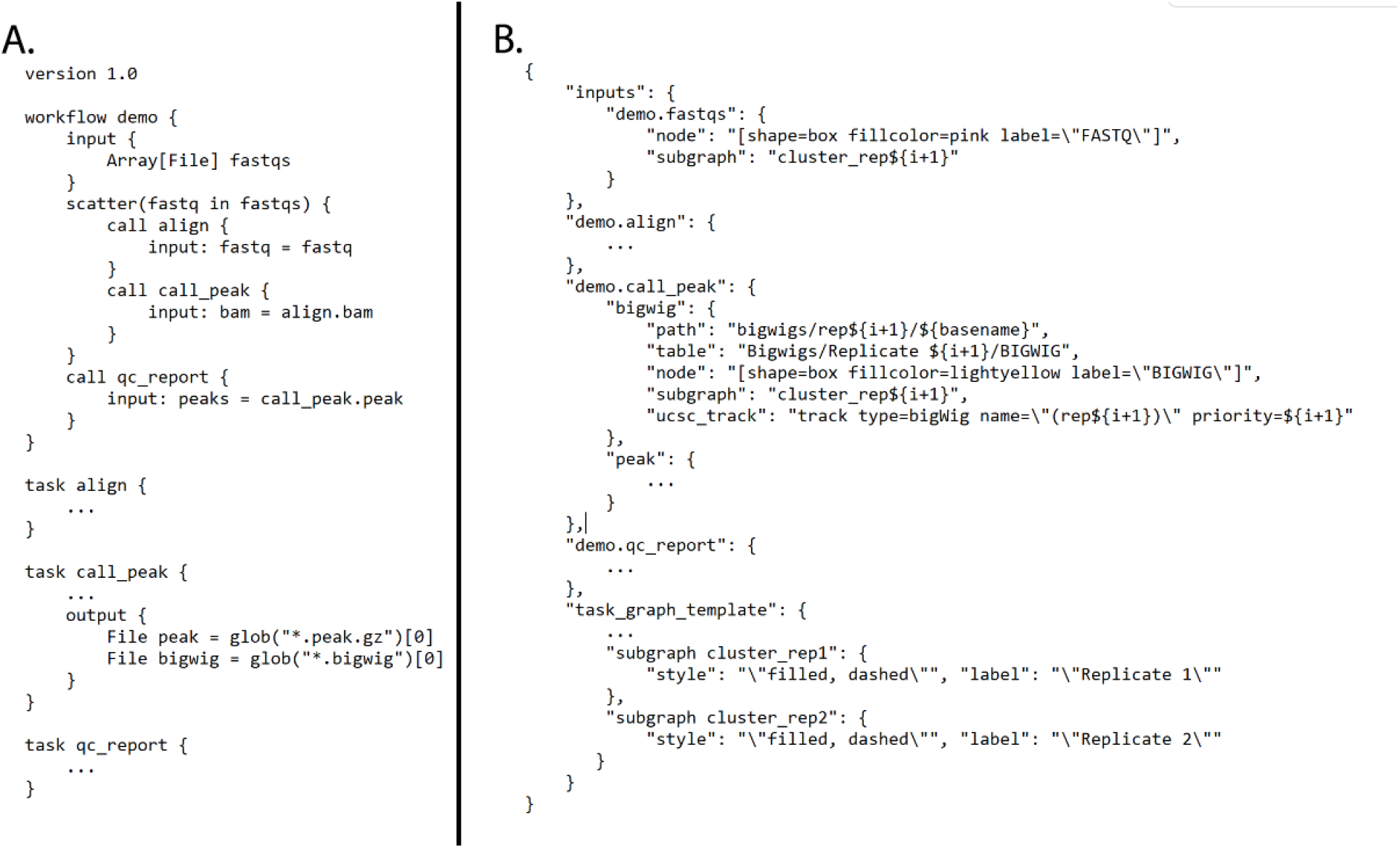
A) Demo WDL pipeline and B) CROO JSON that defines how to organize and display outputs

### Pipeline Infrastructure (CAPER/CROO)

At the scale of a project like ENCODE, the software infrastructure needs infrastructure. Running 2 or 7 or 12 datasets through a pipeline is fairly manageable, but the final phase of ENCODE required us to run 14,000+ datasets (at least 40,000 fastqs) across about 20 different assays, each with its own pipeline and/or set of parameters. To assist us with efficient workflow submissions, we developed the CAPER software package (https://github.com/ENCODE-DCC/caper). CAPER, or “Cromwell-Assisted Pipeline ExecutoR” is a python wrapper for Cromwell, based on UNIX utilities, cloud platform python libraries (google-cloud-storage and boto3) and CLIs (curl, gsutil and aws-cli). It provides a user-friendly terminal based interface to Cromwell by composing the necessary inputs and automatic file transfer between local disks and cloud storage.

CAPER uses a REST API and a mysql/postgresql database to manage Cromwell on a variety of platforms as needed. Typically, a server is instantiated on a machine or cloud instance and is used to marshal input files and parameters (“input.json”) and pass them forward into the WDL]/Cromwell/Docker system. CAPER can localize input files between two different platforms such as Google Cloud Storage (GCS: gs://), AWS S3 (s3://) and a local file system. For example, if input files are provided as S3 URIs and a pipeline is submitted on Google Cloud Platform, then CAPER localizes S3 files on GCS first and passes them to Cromwell.

CROO or “Cromwell Output Organizer” (https://github.com/ENCODE-DCC/croo) is a simple python package that was developed by us to assist people outside of the ENCODE Data Coordination Center (DCC) to find and organize the outputs from the pipelines (Fig 2B). CROO can localize and organize outputs between different platforms similarly to how CAPER does. CROO creates simple HTML interfaces with file tables and connectivity graphs, task graphs and UCSC Genome Browser^15^ tracks (Fig 3). CROO provides an additional feature that allows the generation of pre-signed file URIs on cloud providers enabling visualization of private data with any graphics on genome browsers that can access data *via* URI. This allows public genome browsers to view files that would otherwise be hosted privately. Both CAPER and CROO are registered to PyPI (the Python Package Index) such that they can be installed easily with a single shell command line.

**Figure 3.**
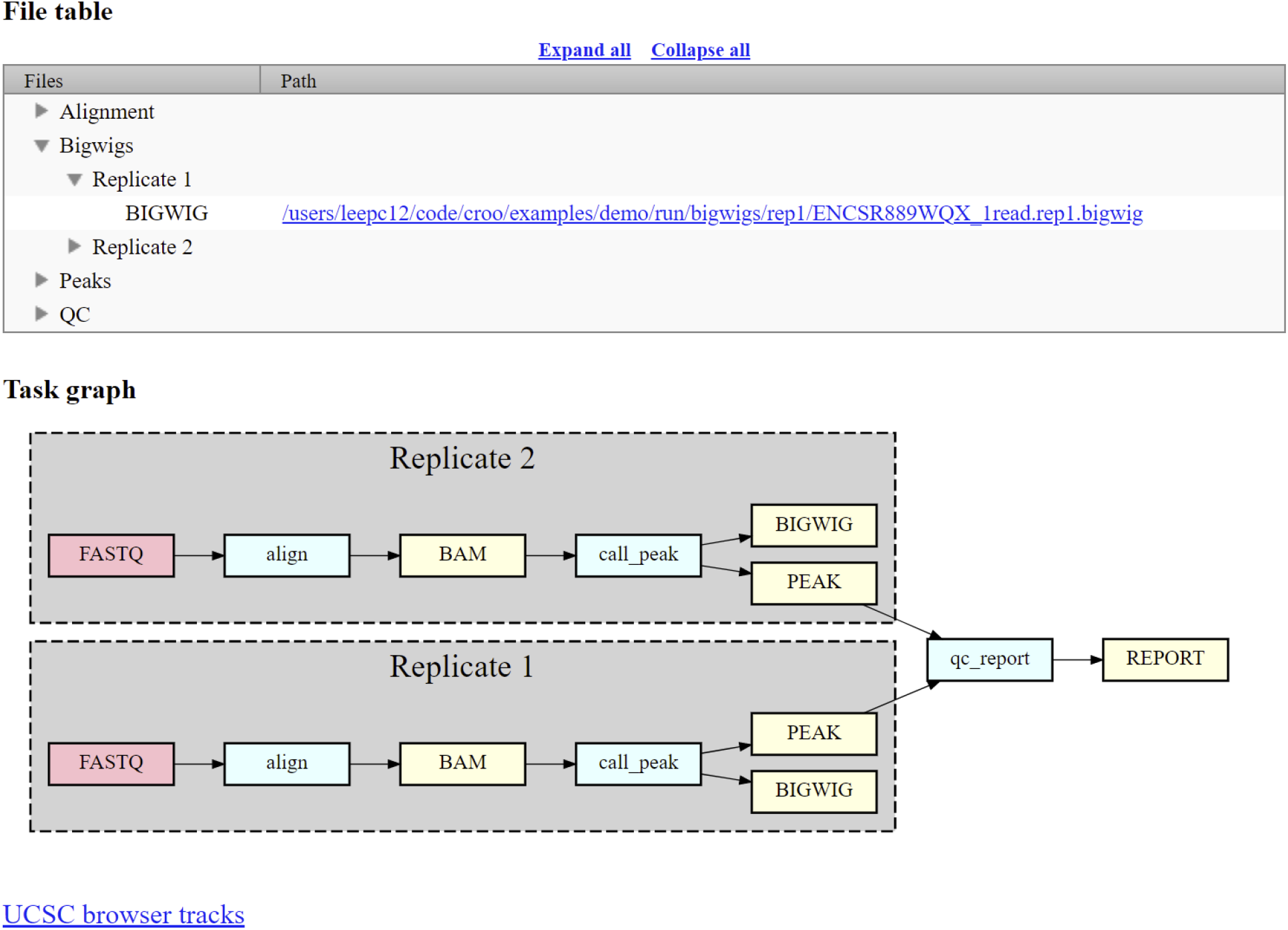
Croo HTML report example showing file table, task graph, and link to UCSC genome browser. The red boxes represent raw data files, the blue boxes represent software steps (abstract names), and the yellow boxes represent intermediate or output processed data files.

### Software and Pipeline Metadata and Provenance

At the DCC itself, we do not use CROO to handle the output of the uniform processing pipelines. In order to carefully track all the provenance, quality metrics, and file relationships required by the ENCODE Portal (Kagda, et al in preparation) we developed a particular data structure that represents each pipeline, quality metric, analysis step, analysis step run, software, and software version. These are all represented in our system as JSON-SCHEMA (https://json-schema.org/) objects in our encodeD instantiation of SnoVault^16^. This pipeline-specific metadata, specifically an object representing an end-to-end analysis, allows us to track the status of runs and create custom pipeline graphs and quality metric reports integrated directly into the ENCODE portal. The common metadata framework we use allows us to integrate results calculated by the DCC using the uniform processing pipelines with any lab- or user-submitted analysis. In effect, we abstract the details of the specific pipeline down to a common framework for visualization and provenance. This allows portal users to have strict confidence in the results that are produced by the consortium. Every output file has a definitive raw data source, a set of software used in every step of its formation - including specific versions of code used to produce this *particular* file, and quality metrics as agreed upon by the consortium.

To map pipeline outputs to the portal we use a custom python package called accession (https://github.com/ENCODE-DCC/accession), which is extended for every official ENCODE uniform processing pipeline. Accession parses the Cromwell workflow metadata and pipeline QC outputs in order to generate the appropriate metadata objects on the ENCODE portal and uploads the data files to the ENCODE AWS S3 bucket. It also supports multiple Cromwell backends (e.g. Google Cloud platform, Amazon EC2, local/HPC) to allow for submission of uniform processing pipeline results from different compute backends.

### The ENCODE ChIP-seq Pipelines

Chromatin-Immunoprecipitation followed by sequencing, or ChIP-seq experiments are at the core of the ENCODE project. This type of assay is used to determine the chromosomal coordinates for binding of transcription factors (TF) and modified histones. We currently house the results of over 5800 ChIP-seq assays from ENCODE in human and mouse, including hundreds of multiplexed MINT-ChIP^7^ modified histone assays. In addition we have over 1600 control ChIP-seq experiments, representing either mock-IP, untreated biosample, input DNA, or “wild-type” (in the case of epitope-tagged constructed) control DNA. All of these experiments are processed through the same ChIP-seq processing pipeline. The TF ChIP-seq pipeline protocol is described in detail in Lee et al “Automated quality control and reproducible peak calling for transcription factor ChIP-seq data", *in preparation* (Fig 4A). ChIP-seq experiments targeting diverse DNA binding proteins and histone marks exhibit inherent high variability of signal-to-noise ratio and number of enriched sites (peaks). Hence, the uniform processing of ChIP-seq results is significantly more complicated than other assays in the ENCODE corpus, since it is necessary to estimate multiple, complementary quality control metrics to carefully compare the signal from mapped reads to controls. Furthermore, the noise inherent in peak-calling of TF ChIP-seq experiments necessitates the use of the Irreproducible Discovery Rate^17^ (IDR) framework to adaptively threshold and retain peaks that are reproducible and rank-concordant across replicates. The latest ENCODE Transcription Factor ChIP (TF-ChIP-seq) pipelines produce, per replicate, two BAM files (filtered and unfiltered alignments), two bigwig files (signal p-value and fold change over control), two peak files (one ranked and one thresholded) and a bigBed file for the IDR thresholded peaks. When there are >1 replicates (usually 2), each pair of replicates is combined to produce another pair of signal files, four peak files (two ranked. two thresholded), and two bigBed files for the IDR thresholded bed files. The histone ChIP pipeline does not use IDR for replicate concordance since peaks of different types of histone marks tend to cover a broad dynamic range of signal-to-noise ratios. Instead, the histone ChIP-seq pipeline just reports a single bed/bigBed pair containing peaks appearing either in both “true” replicates or two pseudo-replicates.

**Figure 4.**
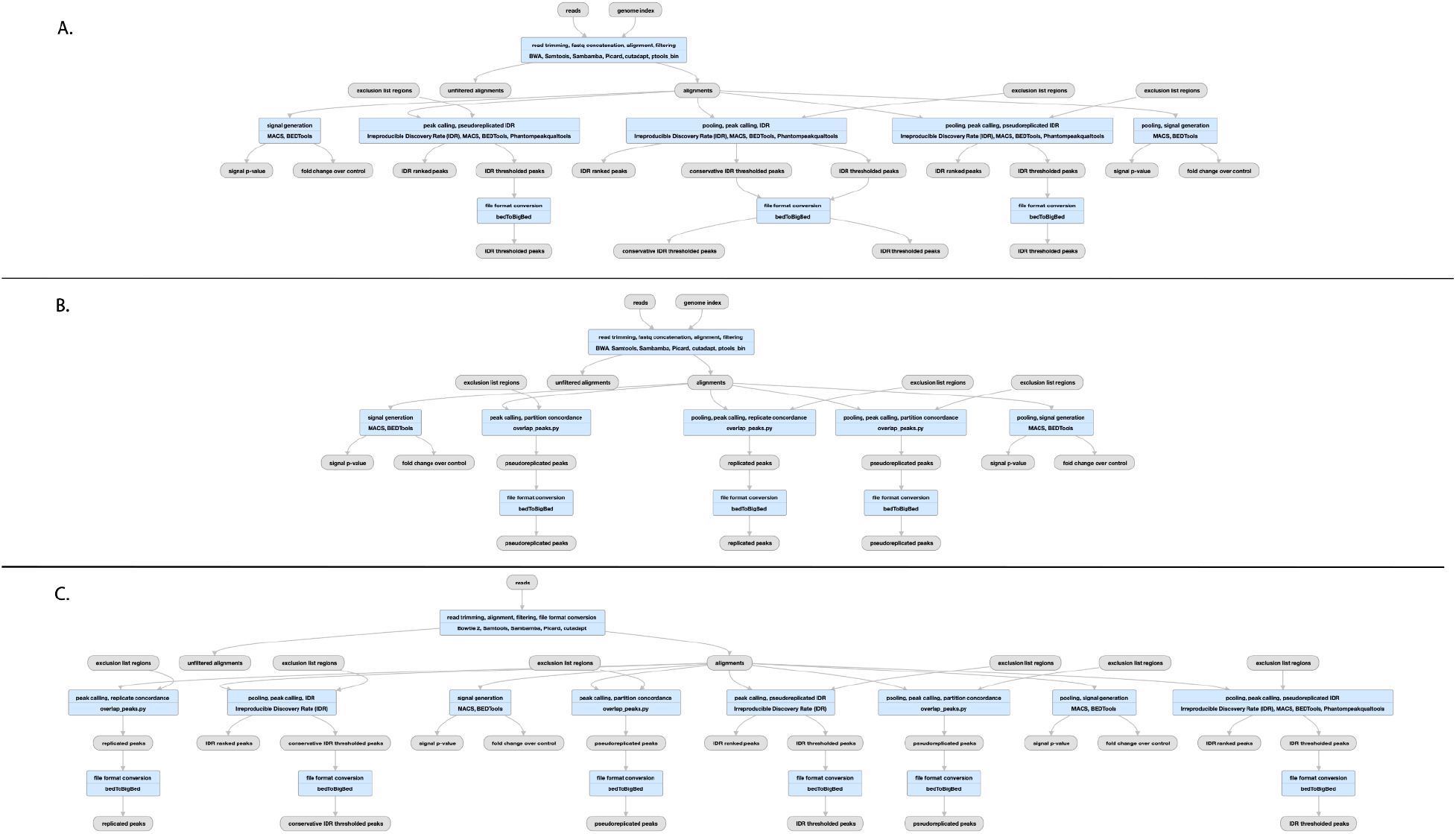
Pipelines for ChIP-seq and ATAC-seq A) TF ChIP-seq schematic; (https://www.encodeproject.org/pipelines/ENCPL367MAS/, B) Histone ChIP-seq schematic; (https://www.encodeproject.org/pipelines/ENCPL612HIG/), ATAC-seq schematic; https://www.encodeproject.org/pipelines/ENCPL787FUN/). Not shown: schematic pipelines for unreplicated experiments; TF ChIP-seq; https://www.encodeproject.org/pipelines/ENCP481MLO/, Histone ChIP-seq; https://www.encodeproject.org/pipelines/ENCPL809GEM/. ATAC-seq : https://www.encodeproject.org/pipelines/ENCPL344QWT/

The pipeline currently uses bowtie2^18^ for mapping TF and Histone ChIP, while the MINT-ChIP experiments use bwa-mem^19^ mapper (Fig 4B). The SPP^20^ peak caller is used to call punctate peaks for TF ChIP-seq experiments, whereas MACS2^21^ is used to call peaks for histone ChIP-seq experiments. The peaks called by the pipeline are filtered utilizing exclusion lists that contain genomic regions resulting in anomalous, unstructured, or experiment independent high signal^22^. Detailed read mapping statistics are used to estimate read quality and mapping rates. The key enrichment QC metrics are “Fraction of Reads In Peaks” (FRIP), normalized and relative strand cross-correlation scores (NSC/RSC)^8^ and Jensen Shannon Distance^23^ metrics between sample and background coverage. Reproducibility of peak calling is estimated using the rescue ratio and self-consistency ratios which compare the number of replicated peaks across and within replicate experiments . Library complexity measurements - the PCR bottleneck coefficients (PBC) and non-redundant fraction (NRF) scores are also calculated.

Thresholds are defined for each of the key quality metrics to assign intuitive levels of potential data quality issues indicated as yellow, orange, or red audit badges on the ENCODE portal. There are actually four slightly different versions of the pipeline, depending on whether the “chipped” factor is a modified histone (https://www.encodeproject.org/pipelines/ENCPL612HIG/, https://www.encodeproject.org/pipelines/ENCPL809GEM/) or transcription factor (https://www.encodeproject.org/pipelines/ENCPL367MAS/, https://www.encodeproject.org/pipelines/ENCPL481MLO/) and whether or not the experiment has replicates.

The performance of the whole pipeline depends on the sequencing depth of the datasets and the size of the genome of interest. Total CPU time ranges from between 1 and 8 hours (average is 2) per million reads and can require up to 18GB of RAM (average is 12 GB).

### The ATAC-seq Pipeline

The ENCODE ATAC-seq pipeline is a small modification of the histone ChIP-seq pipeline (Fig 4C). It uses the same mapper (bowtie2). However, the specific adapters used in the ATAC-seq experiment must be trimmed off prior to mapping to the reference genome.The MACS2 peak caller is used for peak calling with some modifications. One primary difference is that ATAC-seq experiments do not have matched control as a signal baseline. Also, 5’ ends of reads are shifted in a strand-specific manner to account for the Tn5 shift and identify the precise cut-sites. The shifted read-start coverage is aggregated over both strands and smoothed using a 150 bp window for peak calling in MACS2. While IDR is used to estimate reproducibility and stringent peak calls, the default “replicated” peaks are those that are identified by MACS2 with relaxed thresholds in two “true” replicates or two pseudo-replicates. The QC reports for ATAC differ slightly from ChIP-seq, with an emphasis on the Transcription Start Site enrichment score, and the total number of peaks identified.

### The ENCODE RNA-seq Pipelines

The ENCODE (bulk) RNA-seq pipeline (Fig 5A) was developed by the consortium over a period of almost 7 years. It has been used to process data from a menagerie of RNA-seq experiments over the balance of the ENCODE project. Specifically we have processed experiments that have used a wide variety of RNA enrichments, including size (<200 bp), polyadenylation (plus and minus), total, nuclear and other subcellular localizations as well as a series of knockdown quantifications from a variety of methods (siRNA, shRNA, and CRISPRi). The pipeline also works with different library preparation protocols (paired or unpaired reads; with or without strand-specificity). In all cases the pipeline typically produces a common set of files for each replicate: Two BAM files (one each for mapping to the reference genome and transcriptome), three quantifications files (one gene and two transcript; see below) and either two or four signal (bigWig) files. There is one signal file for all reads and one for just uniquely mapping reads, doubled (plus- and minus-strand) if the library is stranded. “Small” RNAs have no transcript quantifications.

**Figure 5.**
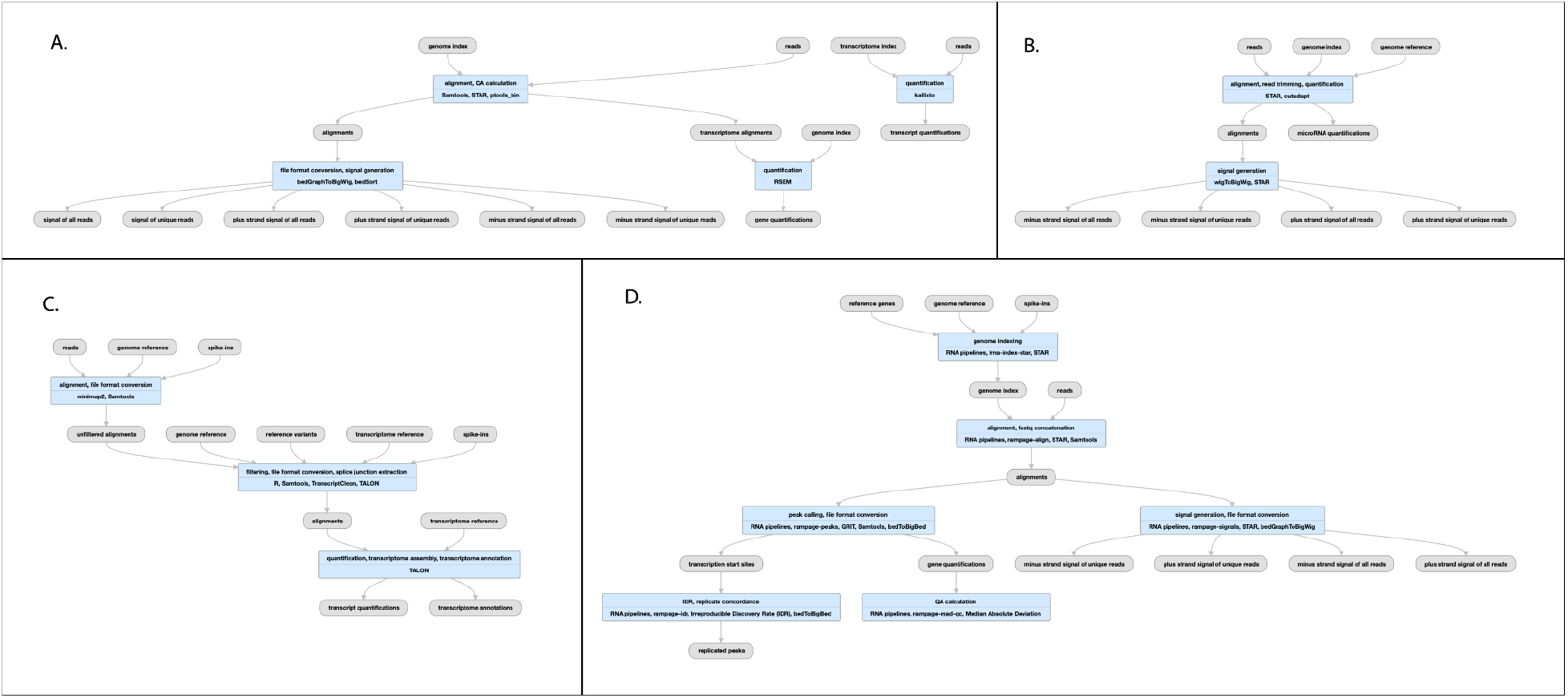
Pipeline for RNA-seq A), bulk RNA seq schematic (https://www.encodeproject.org/pipelines/ENCPL862USL/) B) micro-RNA-seq schematic (https://www.encodeproject.org/pipelines/ENCPL280YDY/) C) long-read RNA-seq schematic (https://www.encodeproject.org/pipelines/ENCPL239OZU/) D) RAMPAGE (and CAGE) schematic (https://www.encodeproject.org/pipelines/ENCPL122WIM)

The core of the pipeline is a mapping or alignment step and a RNA quantification step, with some additional minor steps to process outputs. We use STAR 2.5.1b^24^ to map raw fastq data to both a reference genome (both GRCh38 (https://www.encodeproject.org/files/ENCFF598IDH/) and GRCh37 aka hg19 ([https://www.encodeproject.org/files/ENCFF826ONU/) have been used for human data; GRCm38 aka mm10 has been used for mouse) and reference transcriptome. For transcriptome we have used various versions of GENCODE (https://www.encodeproject.org/files/ENCFF538CQV) including predicted tRNAs. The current versions used in the 4th phase of ENCODE are GENCODE V29 for human and GENCODE M21 for mouse. Older versions of the pipeline also used tophat^25^ for alignment, but this feature was dropped in the current version. For gene and transcript quantification, RSEM^26^ is used to process the BAM files into tsv files that report TPM and FPKM values for all genes and transcripts in the reference annotation (GENCODE) set. For this final phase of the ENCODE project, we added Kallisto^27^ as an alternate, reference-free quantification method, and provide transcript quantifications for both. All the reference files used by the pipeline can also be found at this link: https://www.encodeproject.org/references/ENCSR151GDH

The RNA-seq pipeline implemented for ENCODE produces a variety of QC metrics. In addition to samtools flagstats mapping quality information (https://github.com/samtools) and STAR’s own quality metrics we calculate the number of genes detected and a set of Median Absolute Deviation (MAD) metrics and a plot^28^. We have found that on Google Cloud this pipeline requires about 1 CPU hr/4GB per million reads, with a maximum memory footprint of 120GB.

### micro-RNA

The ENCODE uniform processing microRNA pipeline has been used to process ∼400 datasets submitted from phases 3 and 4 and the REMC project (Fig. 5B). Briefly, Cutadapt^31^ v. 1.7.1 is used to trim the 5’ and 3’ adapters followed by mapping to a transcriptome (GENCODE V29 for human, M21 for mouse) using STAR 2.5.1b to quantify the read counts. The pipeline was modified from that published in^32^ under the direction of the Mortazavi lab. All reference files used for running this pipeline can be found here: https://www.encodeproject.org/references/ENCSR608ULQ

Several QC metrics are calculated for microRNA-seq runs; specifically the mapped read depth, replicate concordance, and number of uRNAs detected. Computational runs use about 0.5 CPU hours and 2 GB/hours per million reads, with a maximum memory footprint of 60GB.

### long read RNA

ENCODE has currently produced approximately 200 long-read RNA-seq data sets in human and mouse from both Pacific Biosciences (PacBio) and Oxford Nanopore (ONT) platforms. These experiments are designed for full-transcript discovery and quantification, and the more standard bulk RNA-seq pipelines are not appropriate for these long reads. Dana Wyman and others in the Mortazavi lab created the TALON (Wyman et al: http://www.biorxiv.org/content/10.1101/672931v2.full) package specifically for the analysis of this data. With their assistance, the ENCODE DCC packaged their software into our Docker/Cromwell/WDL system to uniformly process long-read RNA-seq data (Fig 5C). TALON has six steps. First, Minimap2^29^ is used to align to a genomic reference. Then, TranscriptClean^30^ corrects non-canonical splice junctions, and flags possible internal priming (cryptic poly-A signals) events. The main TALON software then counts splice junctions and quantifies each transcript. Finally, known transcripts are annotated using GENCODE. The primary QC metric used is the number of genes detected, along with the mapping rate. For details on performance, please refer to Wyman et al, but in our cloud runs a job typically takes about 100 CPU hours per 1 million reads (long-read RNA experiments typically range from 0.5M-3.5M reads), and requires 120GB of RAM. All the reference files used for this pipeline can be found here: https://www.encodeproject.org/references/ENCSR925QOG

### RAMPAGE and CAGE

The current phase of ENCODE did not produce any Cap-Analysis Gene Expression (CAGE) or RNA Annotation and Mapping of Promoters for the Analysis of Gene Expression (RAMPAGE) experiments; both methods are used to find transcription start sites. We did uniformly process 289 experiments from ENCODE phase 2 and phase 3 and Genomics of Gene Regulation (GGR; https://www.genome.gov/Funded-Programs-Projects/Genomics-of-Gene-Regulation) projects using a modified version of the STAR pipeline mentioned here (Fig 5D). The reads are mapped in a manner similar to the bulk RNA pipeline, but peaks are called with GRIT^33^ and replicates are merged with IDR. Signal files are created with STAR and bedGraphToBigWig^12^. MAD statistics and plots are also provided for each replicate. The full pipeline source code is available here: https://github.com/ENCODE-DCC/long-rna-seq-pipeline/tree/master/dnanexus/rampage but has not been modified to run with the WDL/Cromwell cloud system.

### The ENCODE DNA Methylation (WGBS) Pipeline

The GemBS^34^ pipeline was designed in the Heath lab to analyze large scale WGBS datasets. The pipeline comprises two parts: 1) Gem3, a high performance read aligner and 2) BScall which is a variant caller specifically designed for bisulfite sequencing (Fig. 6). The two components are combined in a highly efficient, parallelizable, state-of-the-art workflow to allow accurate and fast execution. Since Gem3 can handle large indices, the alignment is performed only on a single composite reference avoiding the two step alignment against the converted and another against unconverted reference. In order to determine the cytosine methylation status, BScall uses a Bayesian model to jointly infer the most likely genotype and methylation levels. The latter is achieved using base error probabilities and under/over conversion rates. For details, please refer to Merkel, et al.

**Figure 6.**
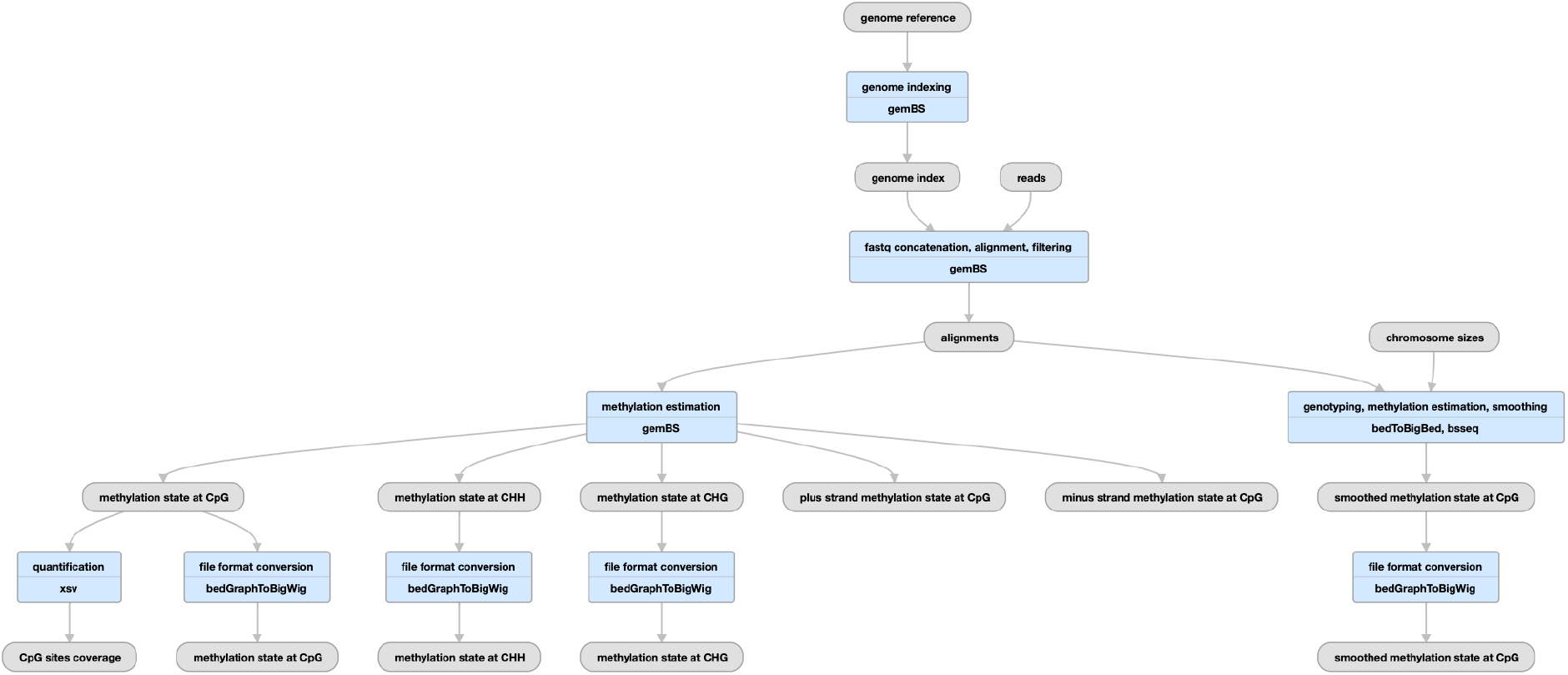
Pipeline schematic using gemBS for whole-genome bisulfite sequencing (https://www.encodeproject.org/pipelines/ENCPL182IUX/)

#### QC metrics

The pipeline produces several useful QC metrics for assessing read mapping, bisulfite conversion efficiency, and replicate concordance. For BAM files, the pipeline computes basic mapping statistics *via* samtools stats (http://www.htslib.org/doc/samtools-stats.html). Using these statistics the pipeline also computes the average coverage for auditing purposes. The pipeline also produces GEM3 mapping quality metrics (http://statgen.cnag.cat/GEMBS/v3/UserGuide/_build/html/qualityControl.html#gem3-report) which includes important WGBS-specific metrics like the lambda conversion rate and general details about mapping efficiency and read quality. For experiments with two replicates, the pipeline calculates the Pearson correlation of the methylation percentage of CpG sites with greater than 10x coverage between the replicates.

These metrics are reflected in the portal metadata, namely in the gemBS alignment quality metrics (https://www.encodeproject.org/profiles/gembs_alignment_quality_metric), CpG correlation quality metrics (https://www.encodeproject.org/profiles/cpg_correlation_quality_metric) and Samtools stats quality metrics (https://www.encodeproject.org/profiles/samtools_stats_quality_metric) which are uploaded to the portal for every pipeline run. Several values in these metrics are automatically checked against the ENCODE standards

A typical execution of the WGBS pipeline takes approximately 0.02 hours (wall time) per million reads based on workflow metadata available on the ENCODE portal. Roughly 70% of this wall time consists of mapping with 16 CPUs and 128 GB of RAM, 14% of the time consists of extracting methylation calls with 16 CPUs and 192 GB of RAM, and 10% of the wall time consists of making methylation and genotype calls using 16 CPUs and 64 GB of RAM. The remaining 6% of wall time consists of preparing configuration files and generating QC statistics and requires significantly less resources.

### The ENCODE DNase-Seq Pipeline

The DNase-seq pipeline has been developed in concert with the Stamatoyannopoulos lab over the past several years (Fig. 7). Initial mapping to the reference genome is performed with BWA^35^, the alignments are filtered and peaks and signal files are created by hotspot2 (https://github.com/Altius/hotspot2). The hotspot software was originally described by John et al.^36^, but numerous improvements have been made in the latest version. hotspot2 counts DNaseI cleavages within a small region ("window") around each site across the genome. It slides this window across the genome, and statistically evaluates cleavage counts within their local context, using a sequence model of DNaseI cleavage sites. The current iteration of the pipeline produces a read-depth normalized signal file (bigWig) and several hypersensitive site peak files (bed and bigBed) thresholded at different false discovery rates (FDR), a genome-wide set of DNaseI cut rates (bed/bigBed) as well as bed/bigBed files for the footprints. For details on the statistics of the footprinting algorithm see the Supplementary Methods of Vierstra et al.^37^

**Figure 7.**
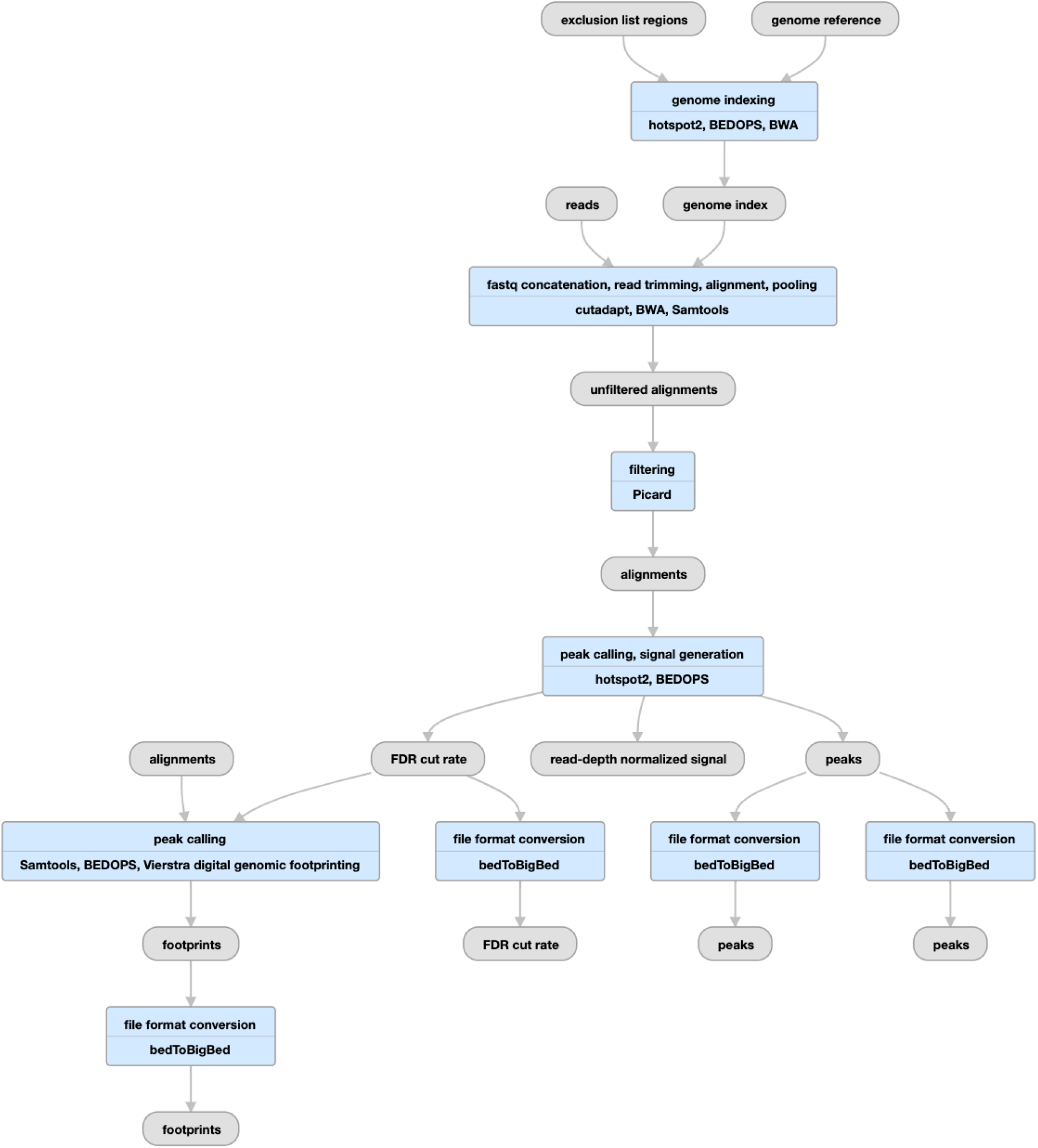
Pipeline schematic for DNase-seq (https://www.encodeproject.org/pipelines/ENCPL848KLD)

Alignment and trimming metrics are calculated by samtools and cutadapt, while other utilities measure the extent of read duplication and fragment size distribution. The key measures used to determine the overall quality of the experiment are the mapped read depth and the SPOT score (“<u>S</u>ignal <u>P</u>ortion <u>O</u>f <u>T</u>ags”). The SPOT score, calculated by hotspot2, is analogous to the FRIP metric used in ATAC-seq and ChIP-seq pipelines. The DNase-seq pipeline on average uses 1.3 hours of CPU time per million reads and has a maximum memory footprint of 32GB.

### The ENCODE Hi-C Pipeline

The ENCODE Hi-C pipeline has been developed with the Aiden lab using their Juicer suite of software tools^38^, with some updates to mapping parameters and chimeric read handling. There are essentially five steps in the pipeline (Fig 8A); mapping (with bwa-mem) and filtering plus Pairix^39^ to form a set of contacts, or pairs file. The genome is then binned into 14 resolutions (between 10bp and 2.5Mbp) by Juicer to form contact matrix (.hic) files. These .hic files can be visualized using Juicebox^40^ or converted to other formats for other visualization software.

**Figure 8:**
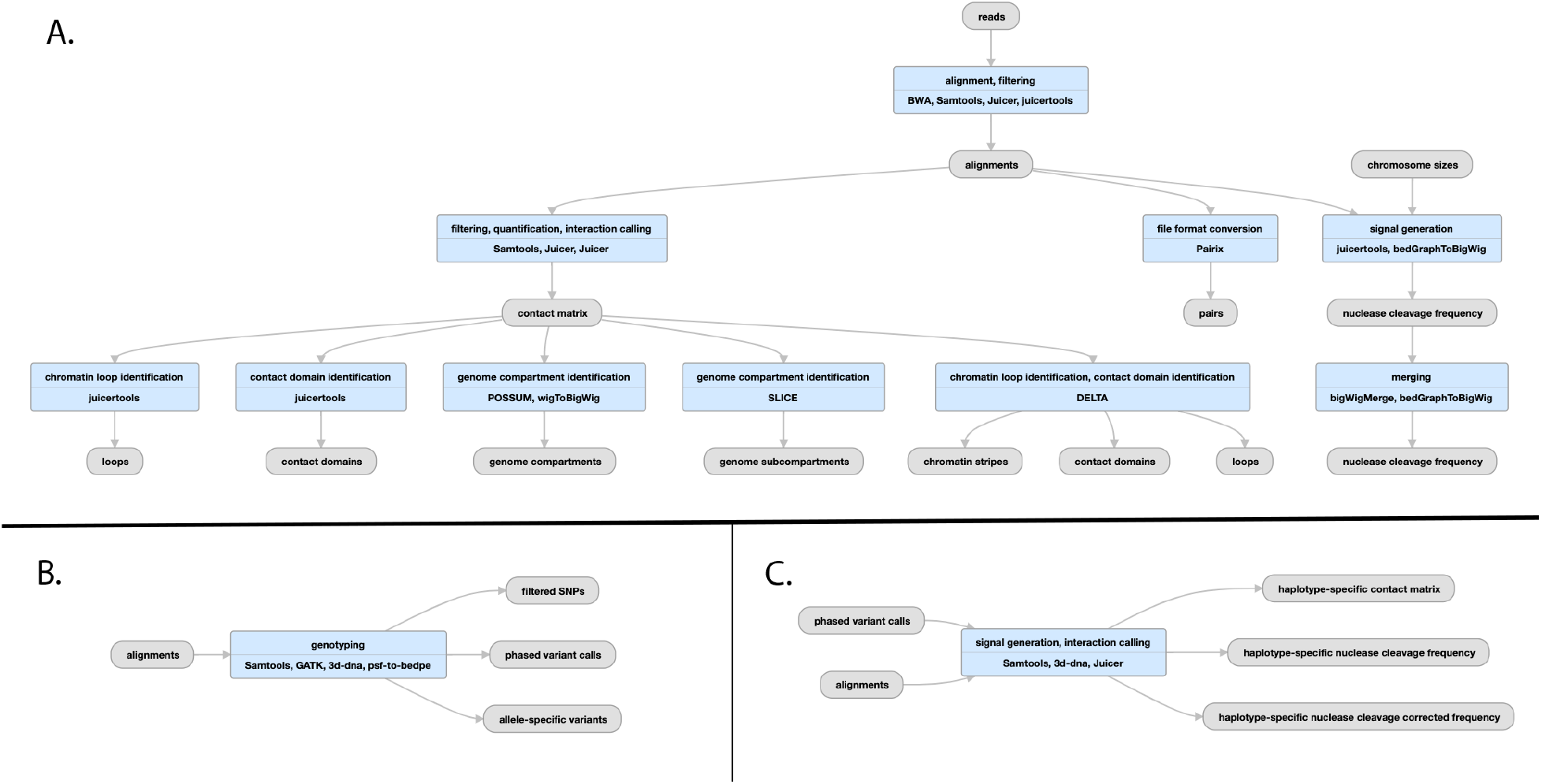
Pipeline schematic for Hi-C pipeline A) Juicer mapping and contact maps schematic: (https://encodeproject.org/pipelines/ENCPL839OAB/). Megamapping is the same but starting from arrays of .hic and .bigWig files merged into deeper maps. B) Genophasing schematic (https://www.encodeproject.org/pipelines/ENCPL780XND/) C) Diploidification schematic (https://www.encodeproject.org/pipelines/ENCPl478DPO/)

HiCCUPS^41^ is used to identify loops while the SLICE and POSSUM utilities identify a/b compartments and subcompartments and the DELTA utility identifies chromatin stripes and contact domains from the contact matrix.

The “diploidification” pipeline comprises two parts: genophase (genotype + phase) and diploidify (Fig. 8B,C). The former experiment is associated with a donor and produces an annotation file set from multiple individual experiments that are derived from the same donor. The second experiment is associated with an individual experiment pertaining to a single donor.

The genophase step calls single nucleotide polymorphisms and attempts to phase them into chromosome-length phased blocks. The SNP are generated from intact Hi-C read alignments by GATK^13^, with slightly modified parameters. The same intact Hi-C data is used to de novo phase SNPs into two haplotypes using the 3D-DNA phasing module^42^. The results are output as a VCF file. In addition to a VCF a variants Hi-C contact matrix and associated bedpe^43^ annotation file are available to help assess the quality of phasing via analyzing the intra-homolog vs inter-homolog contact frequency. The majority of the chromosomes are expected to have most of the SNPs assigned to a haplotype. The overview statistics of phasing performance is included as a Data QA document attached to each genophasing annotation set.

Diploidification uses the largest phased block in the phased VCF file associated with the donor to split individual chromosome data (Hi-C contact map and nuclease cleavage frequency) into two datasets representing different haplotypes. For each chromosome, the two homologous datasets are arbitrarily assigned pseudohaplotype 1 or 2. We do not identify parental haplotypes nor phase across chromosomes; note that assignment of the same pseudohaplotype to different chromosome homologs (chr1, pseudohaplotype 1 and chr2, pseudohaplotype 1) does not imply they indeed belong to the same haplotype and is done for convenience. The pseudohaplotype data is joined to result in two Hi-C contact files and four nuclease cleavage frequency tracks, with and without normalization for SNP density. The chromosome labels are kept the same across the pseudohaplotype files for ease of cross-comparison.

Finally, sets of maps are summed using a megamapping step, creating aggregate maps that enhance contrast and resolution. Sample sets to be aggregated can derive, for instance, from related tissues (such as “left ventricle of heart”, lung, or immune), can reflect a variety of tissues derived from a single individual, or can simply correspond to the collection as a whole.

The pipeline produces QC metrics for bams from individual biological replicates as well as for the contact maps produced by merging data from all biological replicates. The metrics describe in detail the mapping quality, ligation events, and detected Hi-C contacts. In the case of contacts, the QC includes details about long- and short-range interactions, intra- and inter-chromosomal interactions, and more. The full list of available values is described in detail here: https://www.encodeproject.org/profiles/hic_quality_metric

A typical execution of the Hi-C pipeline takes approximately 60 hours of wall time, corresponding to roughly 1.5 CPU hours/million reads. Hi-C, particularly intact Hi-C experiments are quite large (up to 200 billion reads), and some pipeline steps require 512 GB of RAM. CPU time is governed by converting bams to Juicer merged_nodups format (24%), handling chimeric reads (15%), loop calling (13%), initial .hic file creation (11%), deduplication (9%), conversion to 4DN^44^ pairs format (9%), alignment (8%), and contact matrix normalization (8%).

### ENCODE Reference Files

For reproducibility and cross dataset comparisons, it is critical that all experiments from the same organism be mapped to the exact same genome build (and for RNA-seq, the transcriptome as well). Earlier ENCODE experiments were mapped to both hg19 (GRCh37) and GRCh38, but all experiments from the later phase of the project have been solely mapped to GRCh38. All mouse uniform processing, to date, has been on mm10 (GRCm38). The official GENCODE version used by the current phase of ENCODE is V29 for human and M21 for mouse. All references used in uniform- and lab-submitted processings for ENCODE, REMC, modENCODE, MODERN, and GGR are available here: https://www.encodeproject.org/data-standards/reference-sequences (also included are exclusion lists for mapping, spike-ins, tRNAs, and other references used for complete and uniform processing of the ENCODE corpus.

### ENCODE Standards

One of the hallmarks of the decades-long ENCODE project has been its establishment of transparency of genomic assay standards. While the uniform pipelines track thousands of metrics, only a few of them are used to reject or label experiments. Detailed data standards for all experiment types can be found at (https://www.encodeproject.org/data-standards). Audits and badges indicating experiments or files with mild, moderate, or critical issues are summarized at (https://www.encodeproject.org/data-standards/audits/). Further detail about the audit and badge user interface can be found in Davis et al (2018)^45^.

Full reports of all QC metrics for all steps of all pipelines can be found in Supplementary tables 1-6. In addition to scalar metrics, many useful metric plots are available on the ENCODE portal for each analysis run.

### Using or Installing the ENCODE Pipelines

All the pipelines mentioned in this article are open source and can be obtained from GitHub repositories (links below). The tools and the scripts needed for these pipelines have been containerized and pushed automatically to DockerHub, and each pipeline GitHub repository contains the Dockerfile as well as WDL describing the workflow. The pipelines can be run on different platforms including Google cloud and HPC clusters. Since most HPCs do not allow running a Docker container on their compute nodes, Caper provides built-in backends for HPCs such as SGE, SLURM, PBS and LSF to be able to run a pipeline in a Singularity container. We provide Singularity images and a Conda environment installer for several WDL workflows (ChIP and ATAC). This ensures reproducibility of the workflow on multiple platforms.

Several of these pipelines (ChIP-seq, ATAC-seq, RNA-seq, long read RNA-seq, microRNA-seq, WGBS and Hi-C) and their WDL workflows have been deposited to Dockstore (https://dockstore.org/organizations/ENCODEDCC/collections/Pipelines). Dockstore provides an interface to execute the ported pipelines on various platforms (such as DNAnexus (https://dnanexus.com):, Terra^13^, AnVIL^46^). Five of the pipelines (ChIP-seq, ATAC-seq, RNA-seq, long read RNA-seq, and microRNA-seq) have been ported to the Truwl (https://truwl.com/workflows) bioinformatics platform, and two (ChIP-seq and ATAC-seq) are available on the Seven Bridges platform (https://www.sevenbridges.com/platform/)

All of the source code created by the ENCODE DCC is available from GitHub (see Table 1 for individual pipelines):

**Table 1.**
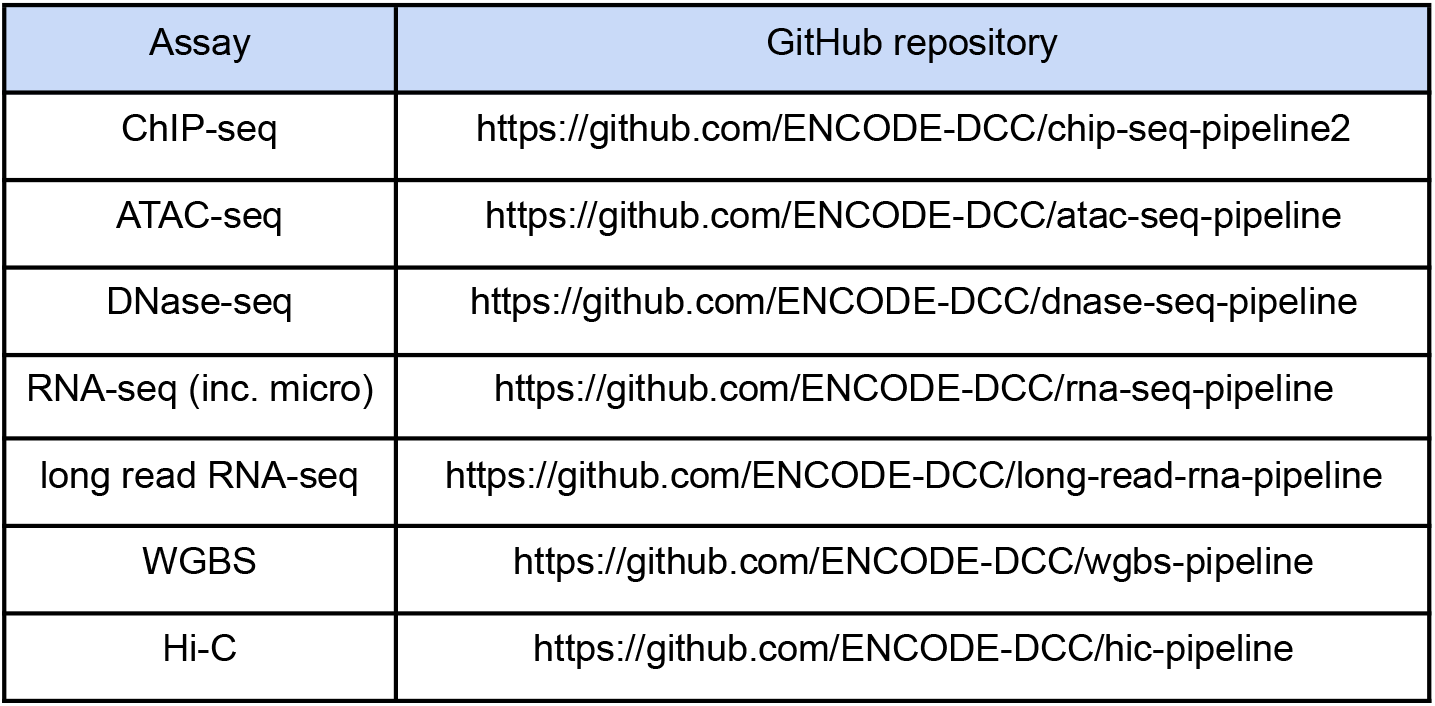
ENCODE DCC implemented uniform processing pipelines.

https://github.com/ENCODE-DCC

https://github.com/ENCODE-DCC/caper

https://github.com/ENCODE-DCC/croo

## Discussion

Much of the information about the uniform processing pipelines at ENCODE can be found at the ENCODE Portal. Each Experiment has a set of processing “frames” called Analyses that constitute a run through the relevant pipeline. Each pipeline execution is captured in the ENCODE metadata with a set of JSON objects representing Analysis Steps, Softwares, Quality Metrics, and most importantly Files (e.g., fastq, bam, bed, bigWig, bigBed, etc.) which are linked to each other with JSON-LD. The inputs (generally starting with fastq files) are connected to the corresponding output files in a graph structure using a “derived_from” pointer-like property that connects files. The graphs for completed runs are presented visually on the ENCODE portal.

Any data file (or other object) that has ever been released publicly remains available to users of the ENCODE portal in perpetuity, although older or deprecated files have a lower status and are not displayed by default.

For the purposes of the ENCODE project, cloud providers such as Google or Amazon have given access to parallel processing power in great excess of our computing needs. We can process or reprocess any arbitrary set of files or experiments, and the “wall clock” time will be equivalent to running a single experiment (on average). Our software and cloud computing APIs make it reasonably straightforward to “spin up” thousands of processors within a few minutes notice.

Developing and maintaining the ENCODE uniform pipelines has been a monumental engineering task. The more experiments that are run through a given pipeline and the more parameters change then more bugs in pipelines and component software will be discovered. In any large-scale effort where thousands of not-necessarily uniform experimental inputs need to be analyzed, users should be prepared to re-run failed jobs as resources are exceeded or parameters need to be adjusted. Since most pipelines are “step-wise”, resources can be saved by restarting pipelines from particular middle points (for example, previously created alignments can be used to re-run the peak calling step). Critical to this endeavor, all pipelines have been created with integrated end-to-end tests, usually wired up to a continuous integration (CI) service. CI runs the tests (usually with a small but complete input dataset) any time a change is pushed to the pipeline github. Even so, as sequencing technologies evolve and as high-throughput sequencing readout experiments get deeper and deeper, failures will occur. One key principle we have striven to uphold is to make all individual pipeline steps idempotent. That is, given the same inputs then the user will always get identical outputs (measured, for example, by equivalent md5 checksums of output files). We caution developers of future bioinformatic pipelines to be judicious in their use of random starting points, or to at least provide a way to input random seeds to their algorithms and software. This ensures that robust engineering of frameworks can be written in a testable manner.

All ENCODE primary and processed data are distributed for free *via* the Amazon Web Services (AWS; https://registry.opendata.aws/encode-project) and the ENCODE portal, https://www.encodeproject.org (a mirror of the data corpus also exists on the Microsoft Azure (https://learn.microsoft.com/en-us/azure/open-datasets/dataset-encode) cloud, courtesy of Microsoft and Terra^13^.

## Acknowledgments

We wish to thank all participants in the ENCODE Consortium for collaborations that have enhanced the metadata definitions, and the ENCODE Portal users for their useful feedback.

## Supplementary Material

[QC description spreadsheets - General.pdf](https://drive.google.com/file/d/1jeN2jvBh5TUTDfnao7vpQ9fMVwVgh4rn/view?usp=share_link)

[QC description spreadsheets - ATAC-seq.pdf](https://drive.google.com/file/d/1h2U5r0BYDIqKmRghjRdgjbf9v2NrkmMw/view)

[QC description spreadsheets - ChIP-seq.pdf](https://drive.google.com/file/d/1_MQZahnJlt-rC5nQMfTtUe1WT4s51Ew6/view)

[QC description spreadsheets - WGBS (gembs).pdf](https://drive.google.com/file/d/1_MQZahnJlt-rC5nQMfTtUe1WT4s51Ew6/view)

[QC description spreadsheets - DNase-seq.pdf](https://drive.google.com/file/d/1EKUiRoE0SXpCEOZhJLwZrZgaU6oLXLRj/view?usp=share_link)

[QC description spreadsheets - RNA-seq (all).pdf](https://drive.google.com/file/d/1i31KhgZ6XjiTa5JdfJupvleD6VFppxOE/view?usp=share_link)

[QC description spreadsheets - Hi-C.pdf](https://drive.google.com/file/d/1ntR_IfnzlSbSHGcWNT878W6Ps5HqGmy3/view)

## References

1. Luo, Y. et al. New developments on the Encyclopedia of DNA Elements (ENCODE) data portal. Nucleic Acids Res. 48, D882–D889 (2020).

2. Jou, J. et al. The ENCODE Portal as an Epigenomics Resource. Curr. Protoc. Bioinformatics 68, e89 (2019).

3. Bernstein, B. E. et al. The NIH Roadmap Epigenomics Mapping Consortium. Nat. Biotechnol. 28, 1045–1048 (2010).

4. Schmidt, D. et al. ChIP-seq: using high-throughput sequencing to discover protein-DNA interactions. Methods 48, 240–248 (2009).

5. Wang, Z., Gerstein, M. & Snyder, M. RNA-Seq: a revolutionary tool for transcriptomics. Nat. Rev. Genet. 10, 57–63 (2009).

6. Boyle, A. P. et al. High-resolution mapping and characterization of open chromatin across the genome. Cell 132, 311–322 (2008).

7. van Galen, P. et al. A Multiplexed System for Quantitative Comparisons of Chromatin Landscapes. Mol. Cell 61, 170–180 (2016).

8. Landt, S. G. et al. ChIP-seq guidelines and practices of the ENCODE and modENCODE consortia. Genome Res. (2012).

9. Batut, P., Dobin, A., Plessy, C., Carninci, P. & Gingeras, T. R. High-fidelity promoter profiling reveals widespread alternative promoter usage and transposon-driven developmental gene expression. Genome Res. 23, 169–180 (2013).

10. Kodzius, R. et al. CAGE: cap analysis of gene expression. Nat. Methods 3, 211–222 (2006).

11. Robinson, P. & Hansen, P. SAM/BAM Format. Computational Exome and Genome doi:10.1201/9781315154770-9/sam-bam-format-peter-robinson-peter-hansen.

12. Kent, W. J., Zweig, A. S., Barber, G., Hinrichs, A. S. & Karolchik, D. BigWig and BigBed: enabling browsing of large distributed datasets. Bioinformatics 26, 2204–2207 (2010).

13. Van der Auwera, G. A. & O’Connor, B. D. *Genomics in the Cloud: Using Docker, GATK, and WDL in Terra*. (‘O’Reilly Media, Inc.’, 2020).

14. Voss, K., Van der Auwera, G. & Gentry, J. Full-stack genomics pipelining with GATK4 + WDL + Cromwell. Preprint at https://doi.org/10.7490/f1000research.1114634.1 (2017).

15. Nassar, L. R. et al. The UCSC Genome Browser database: 2023 update. Nucleic Acids Res. 51, D1188–D1195 (2023).

16. Hitz, B. C. et al. SnoVault and encodeD: A novel object-based storage system and applications to ENCODE metadata. PLoS One 12, e0175310 (2017).

17. Boleu, N., Kundaje, A., Bickel, P. J. & Li, Q. Irreproducible discovery rate. *Berkley, CA*, available at: https://github.com.

18. Langmead, B. & Salzberg, S. L. Fast gapped-read alignment with Bowtie 2. Nat. Methods 9, 357–359 (2012).

19. Li, H. Aligning sequence reads, clone sequences and assembly contigs with BWA-MEM. arXiv [q-bio.GN] (2013).

20. Kharchenko, P. V., Tolstorukov, M. Y. & Park, P. J. Design and analysis of ChIP-seq experiments for DNA-binding proteins. Nat. Biotechnol. (2008).

21. Gaspar, J. M. Improved peak-calling with MACS2. bioRxiv 496521 (2018) doi:10.1101/496521.

22. Amemiya, H. M., Kundaje, A. & Boyle, A. P. The ENCODE Blacklist: Identification of Problematic Regions of the Genome. Sci. Rep. (2019).

23. Ramírez, F. et al. deepTools2: a next generation web server for deep-sequencing data analysis. Nucleic Acids Res. 44, W160–5 (2016).

24. Dobin, A. et al. STAR: ultrafast universal RNA-seq aligner. Bioinformatics 29, 15–21 (2013).

25. Trapnell, C., Pachter, L. & Salzberg, S. L. TopHat: discovering splice junctions with RNA-Seq. Bioinformatics 25, 1105–1111 (2009).

26. Li, B. & Dewey, C. N. RSEM: accurate transcript quantification from RNA-Seq data with or without a reference genome. BMC Bioinformatics 12, 323 (2011).

27. Bray, N. L., Pimentel, H., Melsted, P. & Pachter, L. Erratum: Near-optimal probabilistic RNA-seq quantification. Nat. Biotechnol. 34, 888 (2016).

28. Wu, A. R. et al. Quantitative assessment of single-cell RNA-sequencing methods. Nat. Methods 11, 41–46 (2014).

29. Li, H. Minimap2: pairwise alignment for nucleotide sequences. Bioinformatics 34, 3094–3100 (2018).

30. Wyman, D. & Mortazavi, A. TranscriptClean: variant-aware correction of indels, mismatches and splice junctions in long-read transcripts. Bioinformatics 35, 340–342 (2019).

31. Martin, M. Cutadapt removes adapter sequences from high-throughput sequencing reads. EMBnet.journal 17, 10–12 (2011).

32. Rahmanian, S. et al. Dynamics of microRNA expression during mouse prenatal development. Genome Res. 29, 1900–1909 (2019).

33. Boley, N. et al. Genome-guided transcript assembly by integrative analysis of RNA sequence data. Nat. Biotechnol. 32, 341–346 (2014).

34. Merkel, A. et al. gemBS: high throughput processing for DNA methylation data from bisulfite sequencing. Bioinformatics 35, 737–742 (2019).

35. Li, H. & Durbin, R. Fast and accurate short read alignment with Burrows–Wheeler transform. Bioinformatics 25, 1754–1760 (2009).

36. John, S. et al. Genome-scale mapping of DNase I hypersensitivity. Curr. Protoc. Mol. Biol. **Chapter** 27, Unit 21.27 (2013).

37. Vierstra, J. et al. Global reference mapping of human transcription factor footprints. Nature 583, 729–736 (2020).

38. Durand, N. C. et al. Juicer Provides a One-Click System for Analyzing Loop-Resolution Hi-C Experiments. Cell Syst 3, 95–98 (2016).

39. Lee, S., Bakker, C. R., Vitzthum, C., Alver, B. H. & Park, P. J. Pairs and Pairix: a file format and a tool for efficient storage and retrieval for Hi-C read pairs. Bioinformatics 38, 1729–1731 (2022).

40. Robinson, J. T. et al. Juicebox.js Provides a Cloud-Based Visualization System for Hi-C Data. Cell Syst 6, 256–258.e1 (2018).

41. Rao, S. S. P. et al. A 3D map of the human genome at kilobase resolution reveals principles of chromatin looping. Cell (2014).

42. Hoencamp, C. et al. 3D genomics across the tree of life reveals condensin II as a determinant of architecture type. Science 372, 984–989 (2021).

43. Quinlan, A. R. & Hall, I. M. BEDTools: a flexible suite of utilities for comparing genomic features. Bioinformatics 26, 841–842 (2010).

44. Dekker, J. et al. The 4D nucleome project. Nature vol. 549 219–226 Preprint athttps://doi.org/10.1038/nature23884 (2017).

45. Davis, C. A. et al. The Encyclopedia of DNA elements (ENCODE): data portal update. Nucleic Acids Res. 46, D794–D801 (2018).

46. Schatz, M. C. et al. Inverting the model of genomics data sharing with the NHGRI Genomic Data Science Analysis, Visualization, and Informatics Lab-space. Cell Genom 2, (2022).

