## Supplementary Table 1 for "The ENCODE Uniform Analysis Pipelines"

| SamtoolsFlagstatsQualityMetric | DNase-seq, Total RNA-seq, |
|---|---|
| diff_chroms | flagstats: mate mapped different chr (mapQ>=5) |
| diff_chroms_qc_failed | flagstats: mate mapped different chr (mapQ>=5) - qc failed |
| duplicates | flagstats: duplicates |
| duplicates_qc_failed | flagstats: duplicates - qc failed |
| mapped | flagstats: mapped |
| mapped_pct | flagstats: mapped - percent |
| mapped_qc_failed | flagstats: mapped - qc failed |
| paired | flagstats: paired |
| paired_properly | flagstats: properly paired |
| paired_properly_pct | flagstats: properly paired - percent |
| paired_properly_qc_failed | flagstats: properly paired - qc failed |
| paired_qc_failed | flagstats: paired - qc failed |
| read1 | flagstats: read1 |
| read1_qc_failed | flagstats: read1 - qc failed |
| read2 | flagstats: read2 |
| read2_qc_failed | flagstats: read2 - qc failed |
| singletons | flagstats: singletons |
| singletons_pct | flagstats: singletons - percent |
| singletons_qc_failed | flagstats: singletons - qc failed |
| total | flagstats: total |
| total_qc_failed | flagstats: total - qc failed |
| with_itself | flagstats: with itself and mate mapped |
| with_itself_qc_failed | flagstats: with itself and mate mapped - qc failed |
| usable_fragments | Usable fragments, based on the mapped value. |

| SamtoolsStatsQualityMetric | DNase-seq, WGBS |
|---|---|
| 1st fragments | samtools --stats: 1st fragments |
| average length | samtools --stats: average length |
| average quality | samtools --stats: average quality |
| bases duplicated | samtools --stats: bases duplicated |
| bases mapped | samtools --stats: bases mapped |

| | |
|---|---|
| bases mapped (cigar) | samtools --stats: bases mapped (cigar) |
| bases trimmed | samtools --stats: bases trimmed |
| error rate | samtools --stats: error rate |
| filtered sequences | samtools --stats: filtered sequences |
| insert size average | samtools --stats: insert size - average |
| insert size standard deviation | samtools --stats: insert size - standard deviation |
| inward oriented pairs | samtools --stats: inward oriented pairs |
| is sorted | samtools --stats: is sorted |
| last fragments | samtools --stats: last fragments |
| maximum length | samtools --stats: maximum length |
| mismatches | samtools --stats: mismatches |
| non-primary alignments | samtools --stats: non-primary alignments |
| outward oriented pairs | samtools --stats: outward oriented pairs |
| pairs on different chromosomes | samtools --stats: pairs on different chromosomes |
| pairs with other orientation | samtools --stats: pairs with other orientation |
| raw total sequences | samtools --stats: raw total sequences |
| reads MQ0 | samtools --stats: reads MQ0 |
| reads QC failed | samtools --stats: reads QC failed |
| reads duplicated | samtools --stats: reads duplicated |
| reads mapped | samtools --stats: reads mapped |
| reads mapped and paired | samtools --stats: reads mapped and paired |
| reads paired | samtools --stats: reads paired |
| reads properly paired | samtools --stats: reads properly paired |
| reads unmapped | samtools --stats: reads unmapped |
| sequences | samtools --stats: sequences |
| total length | samtools --stats: total length |
| **CorrelationQualityMetric** | **Deprecated** |
| Pearson correlation | Pearson's R correlation |
| Spearman correlation | Spearman's rank correlation |
| Items | Count of items from two different datasets that are being correlated |

| | |
|---|---|
| Standard deviation | Standard deviation of difference |
| MAD of log ratios | Mean-Average-Deviation (MAD) of replicate log ratios from quantification |
| Details | Description of methods |
