## Supplementary Table 2 for "The ENCODE Uniform Analysis Pipelines"

| HicQualityMetric | |
|---|---|
| sequenced_reads | Number of total reads sequenced for library |
| sequenced_read_pairs | Number of total read pairs sequenced for library |
| no_chimera_found | Number of reads where no chimera was found |
| 0_alignments | Number of reads that did not align to the reference genome |
| 1_alignment | Number of reads that align to one position in the reference genome |
| 1_alignment_unique | Number of unique reads that align to one position in the reference genome |
| 1_alignment_duplicates | Number of duplicate reads that align to one position in the reference genome |
| 2_alignments | Number of reads that align to two positions in the reference genome |
| 2_alignments_a_b | Number of reads that align to two positions in the reference genome of the type A...B |
| 2_alignment_duplicates | Number of duplicate reads that align to two positions in the reference genome |
| 2_alignment_unique | Number of unique reads that align to two positions in the reference genome |
| 2_alignments_a1_a2b_a1b2_b1a2 | Number of reads that align to two positions in the reference genome of the type A1...A2B; A1B2...B1A2 |
| 3_or_more_alignments | Number of reads that align to three or more positions in the reference genome |
| pct_no_chimera_found | Percent of reads where no chimera was found |
| pct_0_alignments | Percent of reads that did not align to the reference genome |
| pct_1_alignment | Percent of reads that align to one position in the reference genome |
| pct_1_alignment_unique | Percent of unique reads that align to one position in the reference genome |
| pct_1_alignment_duplicates | Percent of duplicate reads that align to one position in the reference genome |
| pct_sequenced_1_alignment_unique | Percent of unique reads that align to one position in the reference genome |
| pct_sequenced_1_alignment_duplicates | Percent of duplicate reads that align to one position in the reference genome |
| pct_2_alignments | Percent of reads that align to two positions in the reference genome |
| pct_2_alignment_unique | Percent of unique reads that align to two positions in the reference genome |
| pct_2_alignment_duplicates | Percent of duplicate reads that align to two positions in the reference genome |
| pct_2_alignments_a_b | Percent of reads that align to two positions in the reference genome of the type A...B |
| pct_2_alignments_a1_a2b_a1b2_b1a2 | Percent of reads that align to two positions in the reference genome of the type A1...A2B; A1B2...B1A2 |
| pct_sequenced_2_alignment_unique | Percent of unique reads that align to two positions in the reference genome |
| pct_sequenced_2_alignment_duplicates | Percent of duplicate reads that align to two positions in the reference genome |
| pct_3_or_more_alignments | Percent of reads that align to three or more positions in the reference genome |
| pct_unique_total_duplicates | Percent of duplicate reads that align to the reference genome |
| pct_unique_total_unique | Percent of unique reads that align to the reference genome |
| one_or_both_reads_unmapped | Number of read pairs where one or both reads is unmapped |

| | |
|---|---|
| pct_one_or_both_reads_unmapped | Percent of read pairs where one or both reads is unmapped |
| ligation_motif_present | Number of reads containing a ligation motif |
| pct_ligation_motif_present | Percent of reads containing a ligation motif |
| avg_insert_size | Average insert size |
| total_unique | Number of unique reads that align to the reference genome |
| pct_sequenced_total_unique | Percent of unique reads that align to the reference genome |
| total_duplicates | Number of duplicate reads that align to the reference genome |
| pct_sequenced_total_duplicates | Percent of duplicate reads that align to the reference genome |
| library_complexity_estimate | Estimate of Library Complexity |
| library_complexity_estimate_1_alignment | Estimate of Library Complexity (for reads with one alignment) |
| library_complexity_estimate_2_alignments | Estimate of Library Complexity (for reads with two alignments) |
| library_complexity_estimate_1_and_2_alignments | Estimate of Library Complexity (for reads with one and two alignments) |
| intra_fragment_reads | Number of intra-fragment reads |
| pct_sequenced_intra_fragment_reads | Percent of intra-fragment reads |
| pct_unique_intra_fragment_reads | Percent of unique intra-fragment reads |
| below_mapq_threshold | Number of reads below the mapQ threshold |
| pct_sequenced_below_mapq_threshold | Percent of reads below the mapQ threshold |
| pct_unique_below_mapq_threshold | Percent of unique reads below the mapQ threshold |
| hic_contacts | Number of Hi-C contacts |
| pct_sequenced_hic_contacts | Percent of Hi-C contacts |
| pct_unique_hic_contacts | Percent of unique Hi-C contacts |
| pct_5_prime_bias_long_range | Percent of read ends from long range contacts mapping closer to the 5’ end of the fragment |
| pct_3_prime_bias_long_range | Percent of read ends from long range contacts mapping closer to the 3’ end of the fragment |
| lior_convergence | Distance in basepairs at which left, right, inner, and outer pair types converge |
| pct_left_pair_type | Percentage of Hi-C contacts with left pair type |
| pct_right_pair_type | Percentage of Hi-C contacts with right pair type |
| pct_inner_pair_type | Percentage of Hi-C contacts with inner pair type |
| pct_outer_pair_type | Percentage of Hi-C contacts with outer pair type |
| inter_chromosomal | Number of inter-chromosomal Hi-C contacts |
| pct_sequenced_inter_chromosomal | Percent of inter-chromosomal Hi-C contacts |
| pct_unique_inter_chromosomal | Percent of unique inter-chromosomal Hi-C contacts |
| intra_chromosomal | Number of intra-chromosomal Hi-C contacts |

| | |
|---|---|
| pct_sequenced_intra_chromosomal | Percent of intra-chromosomal Hi-C contacts |
| pct_unique_intra_chromosomal | Percent of unique intra-chromosomal Hi-C contacts |
| short_range_less_than_500bp | Number of intra-chromosomal Hi-C contacts with loci less than 500 basepairs apart |
| pct_sequenced_short_range_less_than_500bp | Percent of intra-chromosomal Hi-C contacts with loci less than 500 basepairs apart |
| pct_unique_short_range_less_than_500bp | Percent of unique intra-chromosomal Hi-C contacts with loci less than 500 basepairs apart |
| short_range_500bp_to_5kb | Number of intra-chromosomal Hi-C contacts with loci between 500 basepairs and 5 kilobases apart |
| pct_sequenced_short_range_500bp_to_5kb | Percent of intra-chromosomal Hi-C contacts with loci between 500 basepairs and 5 kilobases apart |
| pct_unique_short_range_500bp_to_5kb | Percent of unique intra-chromosomal Hi-C contacts with loci between 500 basepairs and 5 kilobases apart |
| short_range_5kb_to_20kb | Number of intra-chromosomal Hi-C contacts with loci between 5 and 20 kilobases apart |
| pct_sequenced_short_range_5kb_to_20kb | Percent of intra-chromosomal Hi-C contacts with loci between 5 and 20 kilobases apart |
| pct_unique_short_range_5kb_to_20kb | Percent of unique intra-chromosomal Hi-C contacts with loci between 5 and 20 kilobases apart |
| long_range_greater_than_20kb | Number of intra-chromosomal Hi-C contacts with loci greater than 20 kilobases apart |
| pct_sequenced_long_range_greater_than_20kb | Percent of intra-chromosomal Hi-C contacts with loci greater than 20 kilobases apart |
| pct_unique_long_range_greater_than_20kb | Percent of unique intra-chromosomal Hi-C contacts with loci greater than 20 kilobases apart |
