## Supplementary Table 3 for "The ENCODE Uniform Analysis Pipelines"

| StarQualityMetric | total RNA-seq, microRNA-seq |
|---|---|
| % of chimeric reads | STAR % of chimeric reads |
| % of reads mapped to multiple loci | STAR % of reads mapped to multiple loci |
| % of reads mapped to too many loci | STAR % of reads mapped to too many loci |
| % of reads unmapped: other | STAR % of reads unmapped: other |
| % of reads unmapped: too many mismatches | STAR % of reads unmapped: too many mismatches |
| % of reads unmapped: too short | STAR % of reads unmapped: too short |
| Average input read length | STAR Average input read length |
| Average mapped length | STAR Average mapped length |
| Deletion average length | STAR Deletion average length |
| Deletion rate per base | STAR Deletion rate per base |
| Insertion average length | STAR Insertion average length |
| Insertion rate per base | STAR Insertion rate per base |
| Mapping speed, Million of reads per hour | STAR Mapping speed, Million of reads per hour |
| Mismatch rate per base, % | STAR Mismatch rate per base, % |
| Number of chimeric reads | STAR Number of chimeric reads |
| Number of input reads | STAR Number of input reads |
| Number of reads mapped to multiple loci | STAR Number of reads mapped to multiple loci |
| Number of reads mapped to too many loci | STAR Number of reads mapped to too many loci |
| Number of splices: AT/AC | STAR Number of splices: AT/AC |
| Number of splices: Annotated (sjdb) | STAR Number of splices: Annotated (sjdb) |
| Number of splices: GC/AG | STAR Number of splices: GC/AG |
| Number of splices: GT/AG | STAR Number of splices: GT/AG |
| Number of splices: Non-canonical | STAR Number of splices: Non-canonical |
| Number of splices: Total | STAR Number of splices: Total |
| Uniquely mapped reads % | STAR Uniquely mapped reads % |
| Uniquely mapped reads number | STAR Uniquely mapped reads number |
| read_depth | Sum of the uniquely mapped reads number and the number of reads mapped to multiple loci. |

| GeneQuantificationQualityMetric | total RNA-seq |
|---|---|
| number_of_genes_detected | Number of Genes Detected |

| GeneTypeQuantificationQualityMetric | total RNA-seq |
|---|---|
| Mt_rRNA | Number of reads assigned to transcripts from the \"Mt_rRNA\" GENCODE biotype; mitochondrial rRNAs |
| antisense | Number of reads in transcripts that overlap the genomic span (i.e. exon or introns) of a protein-coding locus on the opposite strand. |
| miRNA | Number of reads assigned to transcripts from the \"miRNA\" GENCODE biotype; microRNAs |

| | |
|---|---|
| processed_transcript | Number of reads mapped to genomic regions which don't contain an ORF. |
| protein_coding | Number of reads assigned to transcripts from the \"protein_coding\" GENCODE biotype; contain ORFs |
| rRNA | Number of reads assigned to transcripts from the \"rRNA\" GENCODE biotype; encode for rRNAs |
| ribozyme | Number of reads assigned to transcripts from the \"ribozyme\" GENCODE biotype; encode for ribozymes |
| sRNA | Number of reads assigned to transcripts from the \"sRNA\" GENCODE biotype; encode for sRNAs |
| scaRNA | Number of reads assigned to transcripts from the \"scaRNA\" GENCODE biotype; encode for scaRNAs |
| sense_intronic | Number of reads in long non-coding transcript in introns of a coding gene that does not overlap any exons. |
| sense_overlapping | Number of reads in long non-coding transcript that contains a coding gene in its intron on the same strand. |
| snRNA | Number of reads assigned to transcripts from the \"snRNA\" GENCODE biotype; encode for snRNAs |
| snoRNA | Number of reads assigned to transcripts from the \"snoRNA\" GENCODE biotype; encode for snoRNAs |
| spikein | Number of reads assigned to transcripts from the spike ins |
| **MadQualityMetric** | **total RNA-seq** |
| SD of log ratios | Standard Deviation of replicate log ratios from quantification |
| Pearson correlation | Pearson correlation coefficient of replicates from quantification |
| Spearman correlation | Spearman correlation coefficient of replicates from quantification |
| MAD of log ratios | Mean-Average-Deviation (MAD) of replicate log ratios from quantification |
| **MicroRnaMappingQualityMetric** | **microRNA-seq** |
| aligned_reads | Number of aligned reads |
| **MicroRnaQuantificationQualityMetric** | **microRNA-seq** |
| expressed_mirnas | Number of miRNAs expressed |
| **LongReadRnaMappingQualityMetric** | **long read RNA-seq** |
| full_length_non_chimeric_read_count | Quantity of reads that are full-length and do not contain a chimeric adaptor arrangement. |
| mapping_rate | Proportion of reads mapping to the genome |
| **LongReadRnaQuantificationQualityMetric** | **long read RNA-seq** |
| genes_detected | Number of GENCODE genes detected for the replicate |
