## Supplementary Table 4 for "The ENCODE Uniform Analysis Pipelines"

| **DnaseAlignmentQualityMetric** | |
|---|---|
| fourier_transform_eleven | Fourier transform of the insert size at 11 bases |
| insert_size_histogram | Insert size histogram for all reads |
| insert_size_metric | Insert size metric file |
| large_small_ratio | Ratio of long to short reads |
| nuclear_preseq | Total read metrics file |
| nuclear_preseq_targets | Sequencing depth file |
| **DnaseFootprintingQualityMetric** | |
| footprint_count | Total number of DNaseI footprints |
| **DuplicatesQualityMetric** | |
| Reads Examined | Total number of paired and unpaired reads examined |
| Read Duplicates | Total number of paired and unpaired read duplicates |
| UMI Read Duplicates | Number of UMI flagged read duplicates |
| Percent Duplication | Percent of reads that are duplicates |
| Read Pairs Examined | Number of read pairs examined by picard |
| Read Pair Duplicates | Number of read pairs detected as duplicates by picard |
| Read Pair Optical Duplicates | Number of read pairs detected as optical duplicates by picard |
| Unmapped Reads | Number of unmapped detected by picard |
| Unpaired Reads Examined | Number of unpaired reads examined by picard |
| Unpaired Read Duplicates | Number of unpaired reads detected as duplicates by picard |
| Estimated Library Size | Library size in reads as estimated by picard |
| **HotspotQualityMetric** | |
| spot1_score | SPOT score as calculated by Hotspot1 |
| spot2_score | SPOT score as calculated by Hotspot2 |
| hotspot_count | Count of hotspots discovered by Hotspot2 |
| peaks_count | Count of peaks discovered by Hotspot2 |
| total_tags | Count of read tags provided to Hotspot |
| hotspot_tags | Count of read tags discovered to be in hotspots |
| five_percent_allcalls_count | Count of five percent calls |

| | |
|---|---|
| five_percent_hotspots_count | Count of five percent hotspots |
| five_percent_narrowpeaks_count | Count of five percent narrowpeaks |
| tenth_of_one_percent_narrowpeaks | Count of tenth of one percent narrowpeaks |
| **TrimmingQualityMetric** | |
| PE read-pairs processed | Total number read-pairs processed |
| PE read-pairs trimmed | Total number read-pairs trimmed |
| read_1_with_adapter | Number of read1 with adapter |
| read_2_with_adapter | Number of read2 with adapter |
| SE reads processed | Total number (single-end) reads processed |
| SE reads trimmed | Total number (single-end) reads trimmed |
| total_read_pairs_processed | Total number read-pairs processed |
| **EdwbamstatsQualityMetric** | **Deprecated** |
| alignedBy | The aligner used |
| isPaired | If alignment is from paired-end reads this will be set to 1 |
| isSortedByTarget | If the bam is sorted by target location, this is set to 1; if sorted by name, this is set to 0 |
| mappedCount | Count of mapped reads |
| readBaseCount | Count of total bases in reads |
| readCount | Count of all reads |
| readSizeMax | Longest read |
| readSizeMean | Mean size of all reads |
| readSizeMin | Shortest read |
| readSizeStd | Standard deviation of read size |
| targetBaseCount | Number of bases covered in target (e.g. on chromosomes) |
| targetSeqCount | Count of target sequences (e.g. chromosomes) |
| u4mReadCount | Number of randomly sampled uniquely mapped items used in complexity calculation |
| u4mUniquePos | Number of unique positions that sampled uniquely mapped reads were mapped to |
| u4mUniqueRatio | Ratio of unique positions to uniquely mapped reads |
| uniqueMappedCount | Count of reads uniquely mapped |

| FilteringQualityMetric | Deprecated |
| --- | --- |
| pre-filter all reads | Count of all reads prior to filtering |
| pre-filter mapped reads | Count of mapped reads prior to filtering |
| post-filter all reads | Count of all reads after filtering |
| post-filter mapped reads | Count of mapped reads after filtering |
