## Supplementary Table 5 for "The ENCODE Uniform Analysis Pipelines"

| CpgCorrelationQualityMetric | |
|---|---|
| CpG pairs | Number of CpG pairs |
| CpG pairs with atleast 10 reads each | CpG pairs with atleast 10 reads each |
| Pearson correlation | Pearson's correlation of CpG pairs with at least 10 reads each |
| **GembsAlignmentQualityMetric** | |
| sequenced_reads | Number of sequenced reads |
| unmapped_reads | Number of unmapped reads |
| pct_unmapped_reads | Percentage of unmapped reads |
| correct_pairs | Number of correct pairs |
| general_reads | Number of general reads |
| average_coverage | Average coverage |
| pct_general_reads | Percentage of general reads |
| unique_fragments | Number of unique fragments |
| pct_unique_fragments | Percentage of unique fragments |
| conversion_rate | Bisulfite conversion rate |
| reads_under_conversion_control | Number of reads under conversion control |
| pct_reads_under_conversion_control | Percentage of reads under conversion control |
| reads_over_conversion_control | Number of reads over conversion control |
| pct_reads_over_conversion_control | Percentage of reads over conversion control |
| bisulfite_reads_c2t | Number of bisulfite reads C2T |
| pct_bisulfite_reads_c2t | Percentage of bisulfite reads C2T |
| bisulfite_reads_g2a | Number of bisulfite reads G2A |
| pct_bisulfite_reads_g2a | Percentage of bisulfite reads G2A |
