## Supplementary Table 6 for "The ENCODE Uniform Analysis Pipelines"

| ChipAlignmentEnrichmentQualityMetric | TF, Histone |
|---|---|
| subsampled_reads | Number of reads subsampled from the sample for calculation of all cross-correlation metrics |
| estimated_fragment_len | Fragment length/strandshift. This is the estimated fragment length/strand shift for each dataset as estimated by strand shift cross-correlation analysis |
| corr_estimated_fragment_len | Value of correlation function at estimated fragment length. |
| phantom_peak | Strand shift value at which the phantom (false) peak in cross-correlation is observed. This is typically the read length. |
| corr_phantom_peak | Value of cross-correlation function at strand shift corresponding to phantom_peak (which is typically the read length) |
| argmin_corr | Strand shift corresponding to the minimum value of cross-correlation (min_corr) |
| min_corr | Minimum cross-correlation over a sufficiently wide range of strand shifts (typically -100 bp to ~3 times the expected size of DNA fragments based on sonication and size selection protocols). |
| NSC | Normalized strand cross-correlation = FRAGLEN_CC / MIN_CC. Ratio of strand cross-correlation at estimated fragment length to the minimum cross-correlation over all shifts. |
| RSC | Relative cross-correlation coefficient. Ratio of strand cross-correlation at fragment length and at read length |
| auc | The “area under the curve”, with a maximum value of 0.5. Lower values generally indicate higher and more focal enrichment. |
| syn_auc | The expected area under the curve of a perfectly behaved input sample having the same mean sequencing depth of a given sample. This is useful to put the observed AUC into perspective. |
| x_intercept | The point (on the X-axis) at which the curve is 0. This is approximately the percentage of the genome that was not sequenced in a particular sample. |
| syn_x_intercept | The expected X-intercept of a perfectly behaved input sample having the same mean sequencing depth of a given sample. This is useful to put the observed X-intercept into perspective. |
| elbow_pt | The elbow point attempts to measure the position at which the line turns upward. In practice, this is the point at which the plotted line is furthest from the line from the lower-left to the upper-right corner of the graph (equivalent to a perfect input sample with infinite coverage). The point returned is the position on the X-axis of this elbow point and higher values indicate more enrichment. |
| syn_elbow_pt | The expected elbow point of a perfectly behaved input sample having the same mean sequencing depth of a given sample. This is useful to put the observed elbow point into perspective. |
| jsd | The Jensen-Shannon distance between the replicate and the control. |
| syn_jsd | The Jensen-Shannon distance between a given sample and a perfect input sample with the same coverage depth (i.e., the plot generated from the Poisson probability mass function with lambda equal to the mean coverage in the sample). |
| pct_genome_enrich | The approximate percentage of the genome enriched in signal (e.g., bound by a transcription factor or having a certain histone modification). |
| diff_enrich | The differential enrichment between a given sample and that indicated by --JSDsample at the elbow point. |
| ch_div | The CHANCE divergence between the replicate and the control. |

| ChipAlignmentQualityMetric | TF, Histone |
|---|---|
| total_reads | Number of total reads passing QC" |
| total_reads_qc_failed | Number of total reads failing QC |
| duplicate_reads | Number of reads with duplicates passing QC |
| duplicate_reads_qc_failed | Number of reads with duplicates failing QC |
| mapped_reads | Number of mapped reads passing QC |
| mapped_reads_qc_failed | Number of mapped reads failing QC |
| pct_mapped_reads | Percent of mapped reads passing QC |
| paired_reads | Number of paired reads passing QC |
| paired_reads_qc_failed | Number of paired reads failing QC |
| read1 | Number of read1 reads passing QC |
| read1_qc_failed | Number of read1 reads failing QC |
| read2 | Number of read2 reads passing QC |
| read2_qc_failed | Number of read2 reads failing QC |
| properly_paired_reads | Number of properly paired reads passing QC |
| properly_paired_reads_qc_failed | Number of properly paired reads failing QC |
| pct_properly_paired_reads | Percent of properly paired reads passing QC |
| with_itself | Number of reads with both itself & mate mapped passing QC |
| with_itself_qc_failed | Number of reads with both itself & mate mapped failing QC |
| singletons | Number of singletons (unpaired reads) passing QC |
| singletons_qc_failed | Number of singletons (unpaired reads) failing QC |
| pct_singletons | Percent of singletons (unpaired reads) passing QC |
| diff_chroms | Number of reads with mate mapped to different chromosomes passing QC |
| diff_chroms_qc_failed | Number of reads with mate mapped to different chromosomes failing QC |
| usable_fragments | Usable fragments, based on the mapped_reads value. |

| ChipLibraryQualityMetric | TF, Histone |
|---|---|
| unpaired_reads | Number of unpaired reads before filtering |
| paired_reads | Number of paired reads before duplicate filtering |
| unmapped_reads | Number of unmapped reads before duplicate filtering |
| unpaired_duplicate_reads | Number of unpaired duplicates before duplicate filtering |
| paired_duplicate_reads | Number of paired duplicates before duplicate filtering |
| paired_optical_duplicate_reads | Number of paired optical duplicates before duplicate filtering |
| pct_duplicate_reads | Percent of paired duplicates before duplicate filtering |
| total_fragments | Number of fragments before duplicate filtering |
| distinct_fragments | Number of distinct fragments |
| positions_with_one_read | Number of locations to which exactly one read (pair) maps |
| NRF | Non redundant fraction (indicates library complexity). Number of distinct unique mapping reads (i.e. after removing duplicates) / Total number of reads |
| PBC1 | PCR Bottlenecking coefficient 1 = M1/M_DISTINCT where M1: number of genomic locations where exactly one read maps uniquely, M_DISTINCT: number of distinct genomic locations to which some read maps uniquely |
| PBC2 | PCR Bottlenecking coefficient 2 (indicates library complexity) = M1/M2 where M1: number of genomic locations where only one read maps uniquely and M2: number of genomic locations where 2 reads map uniquely |

| ChipPeakEnrichmentQualityMetric | TF, Histone |
|---|---|
| frip | Fraction of reads in the peak file |
| min_size | Smallest peak width |
| 25_pct | 25th percentile of peak widths |
| 50_pct | 50th percentile of peak widths |
| 75_pct | 75th percentile of peak widths |
| max_size | Largest peak width |
| mean | Mean peak width |

| ChipReplicationQualityMetric | TF, Histone |
|---|---|
| reproducible_peaks | Number of peaks called from one replicate or pooled replicates passing reproducibility test by comparing self-pseudoreplicates or pooled pseudo-replicates. |
| idr_cutoff | Irreproducible Discovery Rate (IDR) threshold used to define reproducible peaks |
| rescue_ratio | max(Np,Nt) / min (Np,Nt); Np: Pooled-pseudoreplicate consistent peaks (comparing two pseudoreplicates generated by subsampling pooled reads from Rep1 and Rep2); Nt: True Replicate consistent peaks (comparing true replicates Rep1 vs Rep2) |
| self_consistency_ratio | max(N1,N2) / min (N1,N2); N1: Replicate 1 self-consistent peaks (comparing two pseudoreplicates generated by subsampling Rep1 reads); N2: same as N1 for Rep2 |
| reproducibility | Reproducibility test result for this experiment (pass/fail) |

| ChipSeqFilterQualityMetric | Deprecated |
|---|---|
| NSC | Normalized strand cross-correlation = FRAGLEN_CC / MIN_CC. Ratio of strand cross-correlation at estimated fragment length to the minimum cross-correlation over all shifts. |
| RSC | Relative cross correlation coefficient. Ratio of strand cross-correlation at fragment length and at read length |
| PBC1 | PCR Bottlenecking coefficient 1 = M1/M_DISTINCT where M1: number of genomic locations where exactly one read maps uniquely, M_DISTINCT: number of distinct genomic locations to which some read maps uniquely |
| PBC2 | PCR Bottlenecking coefficient 2 (indicates library complexity) = M1/M2 where M1: number of genomic locations where only one read maps uniquely and M2: number of genomic locations where 2 reads map uniquely |
| fragment length | Fragment length/strandshift. This is the estimated fragment length/strand shift for each dataset as estimated by strand cross-correlation analysis |
| NRF | Non redundant fraction (indicates library complexity). Number of distinct unique mapping reads (i.e. after removing duplicates) / Total number of reads |

| ComplexityXcorrQualityMetric | Deprecated |
|---|---|
| sample size | Total reads sampled (pairs if applicable) |
| paired-end | Reads are paired-ended |

| | |
|---|---|
| read length | Read length |
| fragment length | Fragment length/strandshift. This is the estimated fragment length/strand shift for each dataset as estimated by strand cross-correlation analysis |
| NRF | Non redundant fraction (indicates library complexity). Distinct Locations Mapped / Sampled Reads |
| PBC1 | PCR Bottlenecking coefficient 1 = Single-read Locations / Distinct Locations |
| PBC2 | PCR Bottlenecking coefficient 2 (indicates library complexity) = Single-read Locations / Multi-read Locations |
| NSC | Normalized strand cross-correlation = FRAGLEN_CC / MIN_CC. Ratio of strand cross-correlation at estimated fragment length to the minimum cross-correlation over all shifts. |
| RSC | Relative cross correlation coefficient. Ratio of strand cross-correlation at fragment length and at read length |
| cross_correlation_plot | Cross-correlation plot |
| **HistoneChipSeqQualityMetric** | **Deprecated** |
| nreads | # of starting reads in the pool (if replicated) or experiment (if unreplicated) |
| nreads_in_peaks | # of reads that fall within peaks. |
| npeak_overlap | # peaks overlapping with true replicate or pooled pseudoreplicate peaks |
| Fp | Fraction reads in replicated/stable narrowPeaks (FRiP) from pooled pseudoreplicates |
| Ft | Fraction reads in replicated/stable narrowPeaks (FRiP) from true replicates |
| F1 | Fraction reads in replicated/stable narrowPeaks (FRiP) from replicate 1 self-pseudoreplicates that pass internal pseudoreplication, when self-pseudoreplication is done on unreplicated experiments. |
| F2 | Fraction reads in replicated/stable narrowPeaks (FRiP) from replicate 2 self-pseudoreplicates |
| frip | Best fraction reads in peaks (FRiP) from peaks |
| **IDRQualityMetric** | **Deprecated** |
| Fp | Fraction reads in IDR peaks (FRiP) from pooled pseudoreplicates |
| Ft | Fraction reads in IDR peaks (FRiP) from true replicates |
| F1 | Fraction reads in peaks (FRiP) from replicate 1 self-pseudoreplicates that pass the internal pseudoreplication IDR threshold, when self-pseudoreplication is done on unreplicated experiments. |
| F2 | Fraction reads in peaks (FRiP) from replicate 2 self-pseudoreplicates |
| Np | Number of peaks from pooled pseudoreplicates |
| Nt | Number of peaks from true replicates |
| N1 | Number of peaks from replicate 1 self-pseudoreplicates that pass the internal pseudoreplication IDR threshold, when self-pseudoreplication is done on unreplicated experiments. |
| N2 | Number of peaks from replicate 2 self-pseudoreplicates |
| IDR_cutoff | IDR cutoff threshold for this experiment |
| self_consistency_ratio | IDR self-consistency ratio for this experiment |
| rescue_ratio | IDR rescue ratio for this experiment |
| reproducibility_test | IDR reproducibility test result for this experiment |
| N_optimal | Number of peaks in the IDR optimal set |
| N_conservative | Number of peaks in the IDR conservative set |
| IDR_plot_true | IDR dispersion plot for true replicates |
| IDR_plot_rep1_pr | IDR dispersion plot for replicate 1 pseudo-replicates |
| IDR_plot_rep2_pr | IDR dispersion plot for replicate 2 pseudo-replicates |
| IDR_plot_pool_pr | IDR dispersion plot for pool pseudo-replicates |
| IDR_parameters_true | IDR run parameters for true replicates |
| IDR_parameters_rep1_pr | IDR run parameters for replicate 1 pseudo-replicates |
| IDR_parameters_rep2_pr | IDR run parameters for replicate 2 pseudo-replicates |
| IDR_parameters_pool_pr | IDR run parameters for pool pseudo-replicates |
| frip | Fraction reads in IDR peaks (FRiP) from optimal peaks |
| **IdrSummaryQualityMetric** | **Deprecated** |
| Final parameter values (mu, sigma, rho, and mix) | IDR: Final parameter values (mu, sigma, rho, and mix) |
| IDR cutoff | IDR: IDR cutoff |
| Initial parameter values (mu, sigma, rho, and mix) | IDR: Initial parameter values (mu, sigma, rho, and mix) |
| Number of peaks passing IDR cutoff | IDR: Number of peaks passing IDR cutoff |
| Number of reported peaks | IDR: Number of reported peaks |
| Percent peaks passing IDR cutoff | IDR: Percent peaks passing IDR cutoff |
| Percent reported peaks | IDR: Percent reported peaks |
