## Supplementary Table 7 for "The ENCODE Uniform Analysis Pipelines"

| **AtacAlignmentQualityMetric** | |
|---|---|
| total_reads | Number of total reads passing QC" |
| total_reads_qc_failed | Number of total reads failing QC |
| duplicate_reads | Number of reads with duplicates passing QC |
| duplicate_reads_qc_failed | Number of reads with duplicates failing QC |
| mapped_reads | Number of mapped reads passing QC |
| mapped_reads_qc_failed | Number of mapped reads failing QC |
| pct_mapped_reads | Percent of mapped reads passing QC |
| paired_reads | Number of paired reads passing QC |
| paired_reads_qc_failed | Number of paired reads failing QC |
| read1 | Number of read1 reads passing QC |
| read1_qc_failed | Number of read1 reads failing QC |
| read2 | Number of read2 reads passing QC |
| read2_qc_failed | Number of read2 reads failing QC |
| properly_paired_reads | Number of properly paired reads passing QC |
| properly_paired_reads_qc_failed | Number of properly paired reads failing QC |
| pct_properly_paired_reads | Percent of properly paired reads passing QC |
| with_itself | Number of reads with both itself & mate mapped passing QC |
| with_itself_qc_failed | Number of reads with both itself & mate mapped failing QC |
| singletons | Number of singletons (unpaired reads) passing QC |
| singletons_qc_failed | Number of singletons (unpaired reads) failing QC |
| pct_singletons | Percent of singletons (unpaired reads) passing QC |
| diff_chroms | Number of reads with mate mapped to different chromosomes passing QC |
| diff_chroms_qc_failed | Number of reads with mate mapped to different chromosomes failing QC |
| non_mito_reads | Number of non-mitochondrial reads |
| mito_reads | Number of mitochondrial reads |
| frac_mito_reads | Fraction of mitochondrial reads |
| frac_reads_in_nfr | Fraction of reads in nucleosome free region (NFR) |
| frac_reads_in_nfr_qc_pass | Whether NFR/mono-nuc reads passes QC |
| frac_reads_in_nfr_qc_reason | Reason for frac_reads_in_nfr_qc_pass |
| nfr_over_mono_nuc_reads | Ratio of reads in nucleosome free region (NFR) to reads in mononucleosome region (Mono-Nuc). |
| nfr_over_mono_nuc_reads_qc_pass | Whether NFR/mono-nuc reads passes QC |
| nfr_over_mono_nuc_reads_qc_reason | Reason for nfr_over_mono_nuc_reads_qc_pass |
| nfr_peak_exists | Whether NFR peak exists |
| mono_nuc_peak_exists | Whether mononucleosome peak exists |
| di_nuc_peak_exists | Whether there are peaks in dinucleosome regions (Di-Nuc). |
| usable_fragments | Usable fragments, based on the mapped_reads value. |
